# The landscape of biomedical research

**DOI:** 10.1101/2023.04.10.536208

**Authors:** Rita González-Márquez, Luca Schmidt, Benjamin M. Schmidt, Philipp Berens, Dmitry Kobak

**Affiliations:** Hertie Institute for AI in Brain Health, University of Tübingen, Germany; Tübingen AI Center, Tübingen, Germany; Nomic AI, New York, USA

## Abstract

The number of publications in biomedicine and life sciences has rapidly grown over the last decades, with over 1.5 million papers now being published every year. This makes it difficult to keep track of new scientific works and to have an overview of the evolution of the field as a whole. Here we present a 2D map of the entire corpus of biomedical literature, and argue that it provides a unique and useful overview of the life sciences research. We based our atlas on the abstract texts of 21 million English articles from the PubMed database. To embed the abstracts into 2D, we used the large language model PubMedBERT, combined with *t*-SNE tailored to handle samples of our size. We used our atlas to study the emergence of the Covid-19 literature, the evolution of the neuroscience discipline, the uptake of machine learning, the distribution of gender imbalance in academic authorship, and the distribution of retracted paper mill articles. Furthermore, we present an interactive web version of our atlas that allows easy exploration and will enable further insights and facilitate future research.

## 1 Introduction

The rate of scientific publishing has been increasing constantly over the past century (Larsen and von Ins, 2010; Bornmann and Mutz, 2015), with over one million articles being currently published every year in biomedicine and life sciences alone. Information about academic publications in these fields is collected in the PubMed database, maintained by the United States National Library of Medicine (pubmed.ncbi.nlm.nih.gov). It now contains over 33 million scientific papers from the last 50 years.

This rapid growth of the biomedical literature makes it difficult to track the evolution of biomedical publishing as a whole. Search engines like PubMed and Google Scholar allow researchers to find specific papers given suitable keywords and to follow the citation networks that these papers are embedded in, yet none of them allows exploration of the biomedical literature landscape from a global perspective. This makes it hard to see how research topics evolve over time, how different fields are related to each other, or how new methods and techniques are adopted in different fields. What is needed to answer such questions, is a bird’s-eye view on the biomedical literature.

In this work we develop an approach that enables all of the above: a global two-dimensional atlas of the biomedical and life science literature which is based on the abstracts of all 21 million English language articles contained in the PubMed database. For simplicity, our map is based on the abstract texts alone, and does not rely on other article parts, such as its main text, figures, or references. To create the map, we embedded the abstracts into two dimensions using the transformer-based large language model PubMedBERT (Gu et al., 2021) combined with the neighbor embedding method *t*-SNE (van der Maaten and Hinton, 2008), adapted to handle samples of this size. Our approach allowed us to create a map with the level of detail exceeding substantially previous works (Boyack et al., 2020; Börner et al., 2012).

We argue that our visualization facilitates exploration of the biomedical literature and can reveal aspects of the data that would not be easily noticed with other analysis methods. We showcase the power of our approach in five examples: we studied (1) the emergence of the Covid-19 literature, (2) the evolution of different subfields of neuroscience, (3) the uptake of machine learning in the life sciences, (4) the distribution of gender imbalance across biomedical fields, and (5) the distribution of retracted paper mill articles. In all cases, we used the embedding to formulate specific hypotheses about the data that were later confirmed by a dedicated statistical analysis of the original high-dimensional dataset.

The resulting map of the biomedical research landscape is publicly available as an interactive web page at https://static.nomic.ai/pubmed.html, developed using the deepscatter library (Nomic AI, 2022). It allows users to navigate the atlas, zoom, and search by article title, journal, and author names, while loading individual scatter points on demand. We envisage that the interactive map will allow further insights into the biomedical literature, beyond the ones we present in this work.

## 2 Results

### 2.1 Two-dimensional atlas allows to explore the PubMed database

We downloaded the complete PubMed database and, after initial filtering (see Methods), were left with 20,687,150 papers with valid English abstracts, the majority of which (99.8%) were published in 1970–2021 (Figure S1). Our goal was to generate a 2D embedding of the abstract texts to facilitate exploration of the data.

To annotate our atlas, we chose a set of 38 labels covering basic life science fields such as ‘virology’ and ‘bio-chemistry’, and medical specialties such as ‘radiology’ and ‘ophthalmology’. We assigned each label to the papers published in journals with the corresponding word in journal titles. For example, all papers published in *Annals of Surgery* were labeled ‘surgery’. As a result, 34.4% of all papers received a label, while the rest remained unlabeled. This method misses papers published in interdisciplinary journals such as *Science* or *Nature*, but labels the core works in each discipline. We chose our labels so that they would cover every region of the two-dimensional space. Therefore, despite only having 34% of the papers labeled, we consider this fraction to be representative of the whole landscape.

To generate a two-dimensional map of the entire PubMed database, we first obtained a 768-dimensional numerical representation of each abstract using PubMed-BERT (Gu et al., 2021), which is a Transformer-based (Vaswani et al., 2017) language model trained on PubMed abstracts and full-text articles from PubMed Central. We then reduced the dimensionality to two using *t*-SNE (van der Maaten and Hinton, 2008).

For the initial step of computing a numerical representation of the abstracts, we evaluated several text processing methods, including bag-of-words representations such as TF-IDF (Jones, 1972) and several other BERT-derived models, including the original BERT (Devlin et al., 2019), SBERT (Reimers and Gurevych, 2019), SciBERT (Beltagy et al., 2019), BioBERT (Lee et al., 2020), SPECTER (Cohan et al., 2020), SimCSE (Gao et al., 2021), and SciNCL (Ostendorff et al., 2022). We chose PubMedBERT because it best grouped papers together in terms of their label, quantified by the *k*-nearest-neighbor (*k*NN) classification accuracy when each label is predicted based on the most frequent label of its 10 nearest neighbors (Table 3). For the PubMedBERT representation, this prediction was correct 69.7% of the time (Table 1). For comparison, TF-IDF, which is simpler and faster to compute, yielded lower *k*NN accuracy (65.2%).

**Table 1:** Quality metrics for the embeddings. Acc.: *k*NN accuracy (*k* = 10) of label prediction. RMSE: root-mean-squared error of *k*NN prediction of publication year. Recall: overlap between *k* nearest neighbors in the 2D embedding and in the high-dimensional space. See Methods for details.

| Data | Dim. | Acc. | RMSE | Recall |
| --- | --- | --- | --- | --- |
| PubMedBERT | 768 | 69.7% | 8.4 | – |
| TF-IDF | 4,679,130 | 65.2% | 8.8 | – |
| $t$-SNE(BERT) | 2 | 62.6% | 10.2 | 6.2% |
| $t$-SNE(TF-IDF) | 2 | 50.6% | 11.2 | 0.7% |
| Chance | – | 4.3% | 12.4 | 0.0% |

**Table 2:** Isolatedness metric for several sets of papers. Fraction of *k*-nearest-neighbors of papers from each corpus that also belong to the same corpus (see Methods). The first four rows show corpora selected based on the abstract text; the last two — based on the journal name.

| | $n$ | BERT | TF-IDF |
| --- | --- | --- | --- |
| Covid-19 | 132,802 | **80.6%** | **76.2%** |
| HIV/AIDS | 308,077 | 63.9% | 62.3% |
| Influenza | 90,575 | 57.9% | 64.1% |
| Meta-analysis | 145,358 | 52.6% | 38.5% |
| Virology | 112,807 | 47.7% | 39.1% |
| Ophthalmology | 144,411 | 47.7% | 43.6% |

**Table 3:** *k*NN accuracy of different BERT-based models. This comparison used a subset of the data (training set size: 990,000 labeled papers; test set size: 10,000 labeled papers). For comparison, the *k*NN accuracy values for the TF-IDF and SVD (*d* = 300) representations measured on the same subset were 61.0% and 54.8% respectively.

|  | Average | [CLS] | [SEP] |
| --- | --- | --- | --- |
| BERT | 57.1% | 50.4% | 53.4% |
| SciBERT | 62.1% | 57.0% | 60.9% |
| BioBERT | 64.0% | 62.7% | 65.0% |
| PubMedBERT | 64.4% | 60.4% | **67.7%** |
| SBERT | 64.5% | 60.7% | 62.2% |
| SPECTER | 64.6% | 63.9% | 64.7% |
| SciNCL | 65.9% | 64.6% | 64.6% |
| SimCSE | 57.0% | 53.2% | 52.1% |

For the second step, we used *t*-SNE with several modifications that allowed us to run it effectively on very large datasets. These modifications included uniform affinities to reduce memory consumption and extended optimization to ensure better convergence (see Methods). With these modifications, *t*-SNE performs better than other neighbor embedding methods such as UMAP (McInnes et al., 2018) in terms of *k*NN accuracy and memory requirements (González-Márquez et al., 2022). The resulting embedding showed good label separation, with *k*NN accuracy in 2D of 62.6%, not much worse than in the 4,679,130-dimensional TF-IDF representation.

We interpret the resulting embedding as the map of the biomedical literature (Figure 1). It showed sensible global organization, with natural sciences mainly located on the left side and medical specialties gathered on the right side; physics- and engineering-related works occupied the bottom-left part (Figures S2, S3). Related disciplines were located next to each other: for example, the biochemistry region was overlapping with chemistry, whereas psychology was merging into psychiatry. A *t*-SNE embedding based on the TF-IDF representation had similar large-scale structure but worse *k*NN accuracy (50.6%; Figure S4).

**Figure 1:**
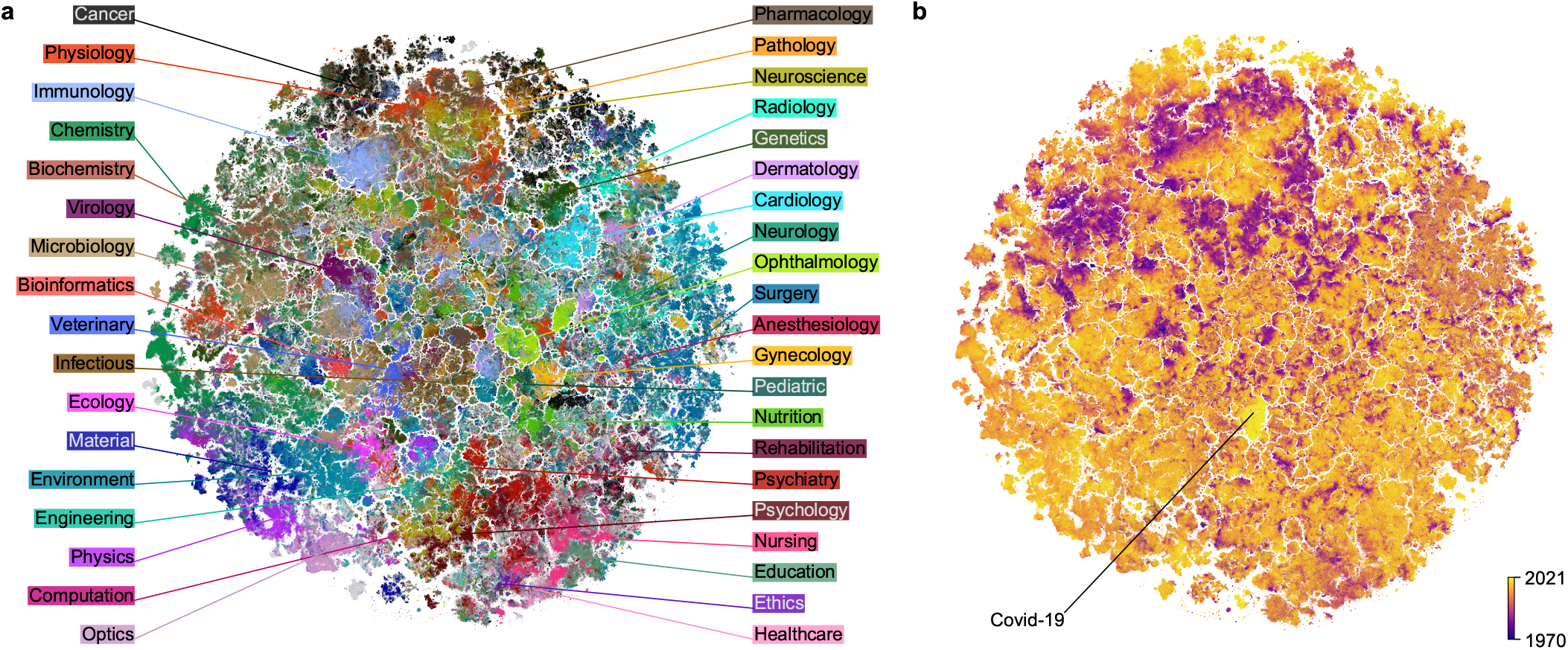
2D embedding of the PubMed dataset. Paper abstracts (*n* = 21 M) were transformed into 768-dimensional vectors with PubMedBERT (Gu et al., 2021) and then embedded in 2D with *t*-SNE (van der Maaten and Hinton, 2008). (a) Colored using labels based on journal titles. Unlabeled papers are shown in gray and are displayed in the background. (b) Colored by publication year (dark: 1970 and earlier; light: 2021).

In addition to this global structure, the map revealed rich and detailed fine structure and was fragmented into small clusters containing hundreds to thousands of papers each (Figure S5a). Even though immediate neighborhoods were distorted compared to the 768-dimensional PubMedBERT representation (only 6.2% of the nearest neighbors in ℝ^2^ were nearest neighbors in ℝ^768^; we call this metric *k*NN recall), manual inspection of the clusters suggested that they consisted of papers on clearly-defined narrow topics.

Moreover, the map had rich temporal structure, with papers of the same age tending to be grouped together (Figure 1b). While this structure may be influenced by changes in writing style and common vocabulary, it is likely primarily caused by research topics evolving over time and becoming more or less fashionable. The most striking example of this effect is a cluster of very recent papers published in 2020–21 that is very visible in the middle of the map (bright yellow in Figure 1b). We will use this island as our first example of how the map can be used to guide understanding of the publishing landscape and how it allows to form hypotheses about the structure and temporal evolution of biomedical research. We will show that these hypotheses can be rigorously confirmed in the high-dimensional embedding space.

### 2.2 The Covid-19 literature is uniquely isolated

The bright yellow island we identified above comprised works related to Covid-19 (Figure 1b), with 85% of papers on Covid-related topics, and 15% on other respiratory epidemics. Our dataset included in total 132,802 Covid-related papers (based on terms such as COVID-19, SARS-CoV-2, etc., present in their abstracts; see Methods), which constituted 5.2% of all PubMed papers published in 2020–2022. As the pandemic and its effects were studied by many different biomedical fields, one might have expected the Covid papers to be distributed across the embedding in their corresponding disciplines. Instead, most (59.3%) of the Covid-related papers were grouped together in one cluster, while the rest were sparsely distributed across the map (Figure S6a).

The main Covid cluster was surrounded by articles on other epidemics, public health issues, and respiratory diseases. When we zoomed in, we found rich inner structure within the Covid cluster itself, with multiple Covid-related topics separated from each other (Figure 2). Papers on mental health and societal impact, on public health and epidemiological control, on immunology and vaccines, on clinical symptoms and treatment — were all largely non-overlapping, and were further divided into even narrower subfields. This suggests that our map can be useful for navigating the literature on the scale of narrow and focused scientific topics.

**Figure 2:**
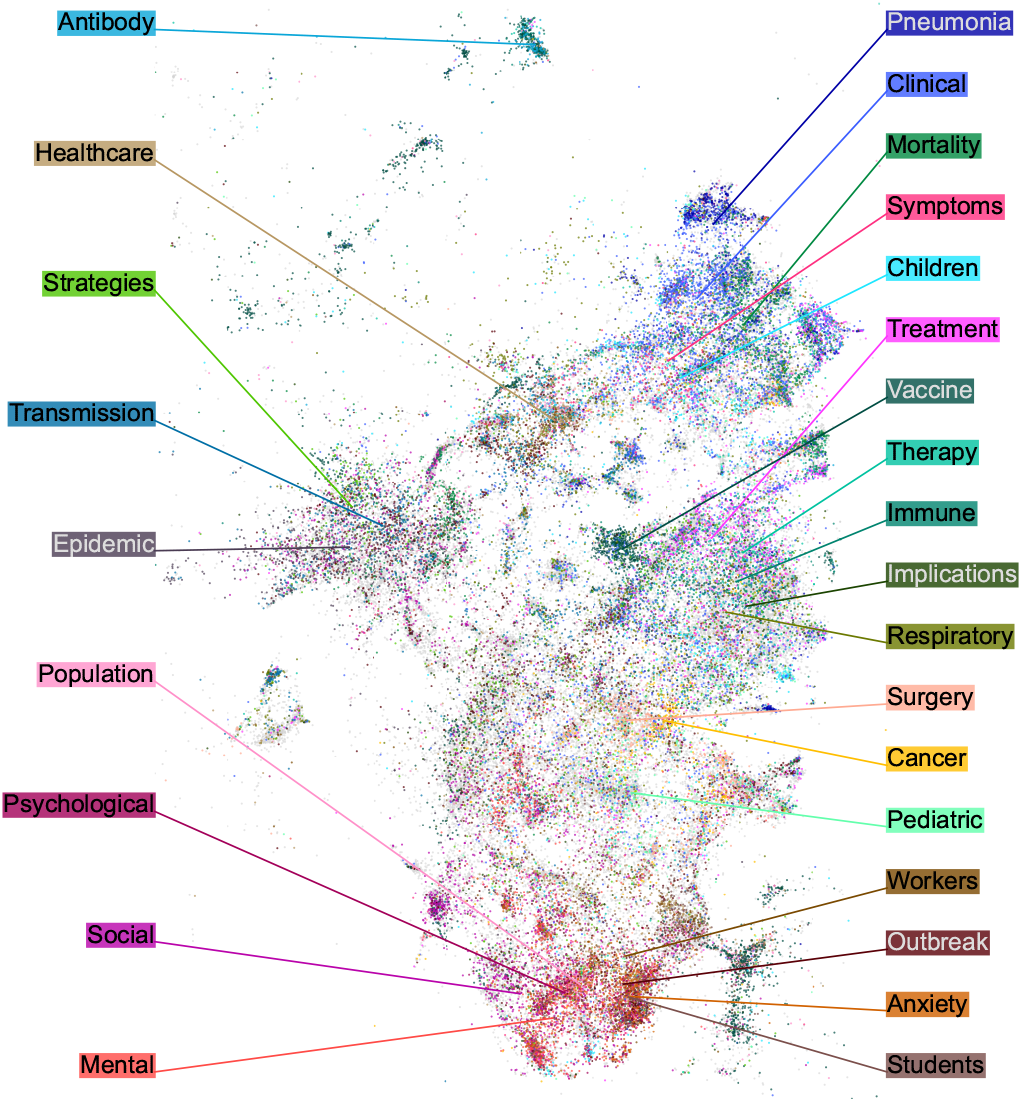
Covid-19 region of the map. Colors are assigned using the most common keywords appearing in paper titles. Uncolored Covid papers are shown in the background in gray. This region in the embedding also contained some non-Covid papers (∼15%) about other respiratory epidemics; they are not shown.

Seeing that the Covid papers prominently stood out in the map (Figure 1b), we hypothesized that the Covid literature was more isolated from the rest of the biomedical literature, compared to other similar fields. To test this, we selected several comparable sets of papers, such as papers on HIV/AIDS or influenza, or all papers published in virology or ophthalmology journals (two labels that appeared particularly compact in Figure 1a). We measured the *isolatedness* of each corpus in the high-dimensional space by the fraction of their *k*NNs that belonged to the same corpus. We found that indeed, Covid literature had the highest isolatedness, in both BERT (80.6%) and TF-IDF (76.2%) representations (Table 2). This suggests that the Covid-19 pandemic had an unprecedented effect on the scientific literature, creating a separate and uniquely detached field of study in only two years.

We investigated the driving factors behind the emergence of the Covid island in the two dimensional space using the TF-IDF representation and saw that, even though the presence of Covid keywords (such as ‘Covid’ or ‘SARS-Cov’) did play some role in the island formation, it was not the only source of similarity between Covid papers (Figure S7).

### 2.3 Changing focus within neuroscience

As we have seen in the extreme example of the Covid literature, the atlas can be used to study composition and temporal trends across disciplines. We next show how it can also provide insights into shifting topics and trends inside a discipline. We demonstrate this using the example of neuroscience. Neuroscience papers (*n* = 240, 135) in the map were divided into two main clusters (Figure 3a). The upper one contained papers on molecular and cellular neuroscience, while the lower one consisted of studies on behavioral and cognitive neuroscience. While it has been shown that articles in brain-related journals show a separation between basic science and clinical applications (Ke, 2019), our map revealed a different bimodality, separating cellular from behavioral neuroscience. Several smaller clusters comprised papers on neurodegenerative diseases and sensory systems.

**Figure 3:**
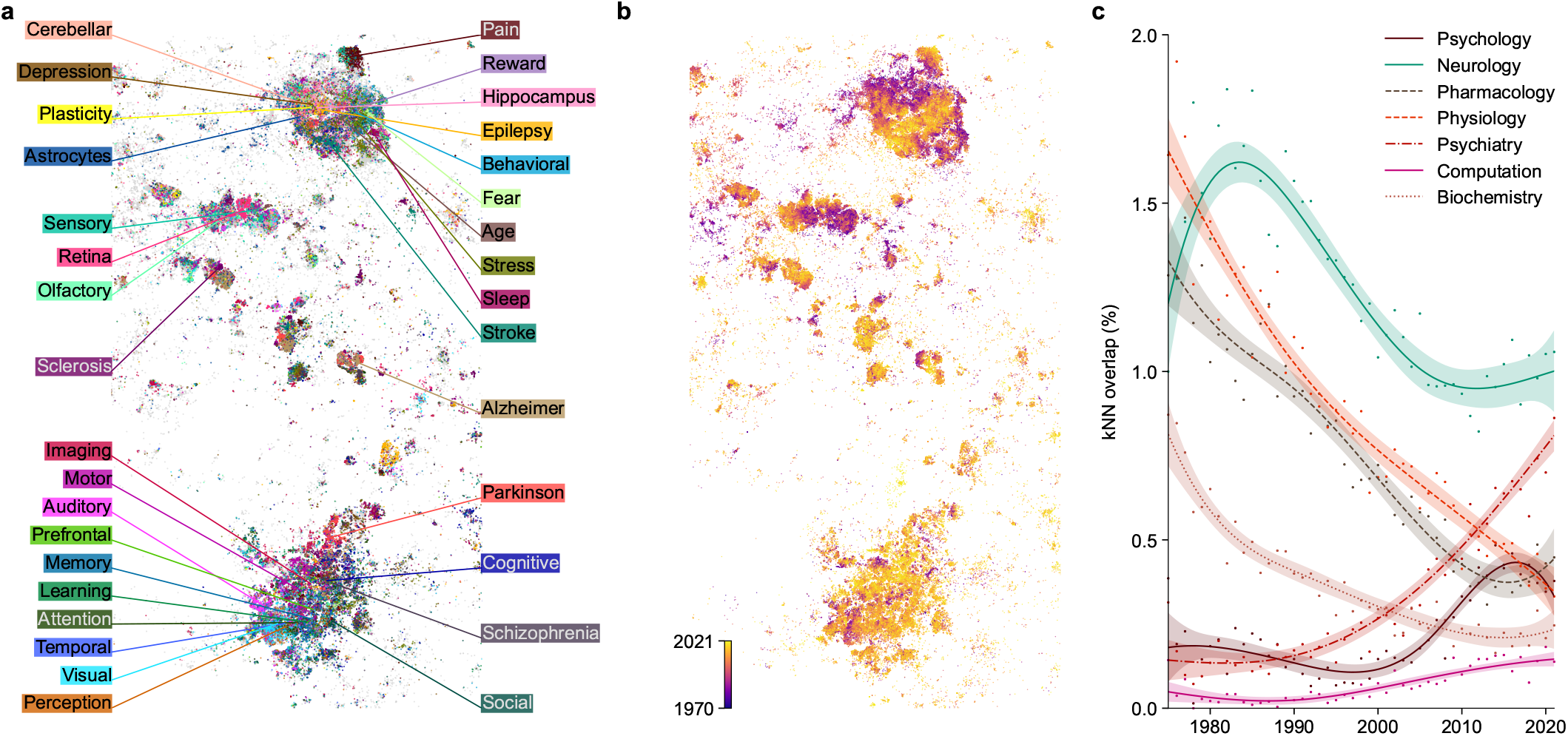
Neuroscience literature. **(a)** Articles published in neuroscience journals, colored by presence of specific keywords in paper titles. **(b)** The same articles colored by the publication year (dark: 1970 and earlier; light: 2021). **(c)** Fraction of the high-dimensional *k*NNs of neuroscience papers that belonged to a given discipline (biochemistry, computation, neurology, pharmacology, physiology, psychiatry, psychology). We chose to analyze those disciplines because they had the highest confusion scores with the neuroscience class in a *k*NN classifier. Points: yearly averages. Smooth curves and 95% confidence intervals were obtained with generalized additive models (see Methods).

Coloring this part of the embedding by publication year indicated that the cellular/molecular region on average had older papers than the cognitive/behavioral region (Figure 3b). This suggests that the relative publication volume in different subfields of neuroscience has changed with time. To test this hypothesis directly, we devised a metric measuring the overlap between neuroscience and any given related discipline across time. We defined *kNN overlap* as the fraction of *k*NNs of neuroscience papers that belonged to a given discipline in the high-dimensional space. We found that the overlap of neuroscience with physiology and pharmacology has decreased since the 1970s, while its overlap with psychiatry, psychology, and computation has increased, in particular after 1990s (Figure 3c). Indeed, neuroscience originated as a study of the nervous system within physiology, but gradually broadened its scope to include cognitive neuroscience, related to psychology, as well as computational neuroscience, related to computer science and machine learning.

### 2.4 The uptake of machine learning

In recent years, computational methods and machine learning have increasingly found use in various biomedical disciplines (Topol, 2019). To explore the use of machine learning (ML) in the biomedical landscape, we computed the fraction of papers claiming to use machine learning (defined as papers mentioning ‘machine learning’ in their abstracts) within different medical disciplines across time (Figure 4a). We found that the uptake of ML differed substantially across disciplines. Radiology was the first discipline to show an increase in ML adoption, shortly after 2015, followed by psychiatry and neurology. In oncology, ML adoption started later but showed accelerated rise over the last five years. This is in contrast with specialties like dermatology and gynecology that did not see any ML usage until ∼2020.

**Figure 4:**
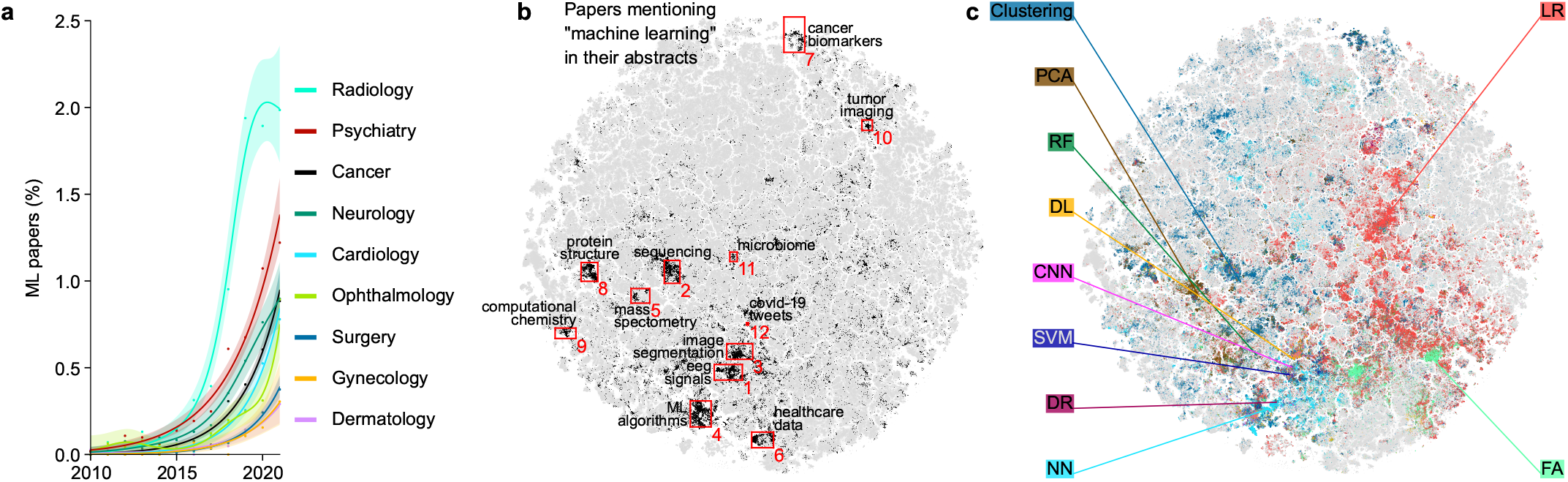
Machine learning papers. **(a)** Percentage of papers mentioning ‘machine learning’ in their abstracts across time for different medical disciplines. Smooth curves and 95% confidence intervals were obtained using generalized additive models and the points correspond to yearly percentages (see Methods). **(b)** Papers mentioning ‘machine learning’ in their abstracts, grouped into 12 clusters that we manually labeled. **(c)** Papers colored according to various statistical and machine learning methods mentioned in their abstracts. Abbreviations: principal component analysis (PCA), random forest (RF), deep learning (DL), convolutional neural network (CNN), support vector machine (SVM), dimensionality reduction (DR), neural networks (NN), linear regression (LR), factor analysis (FA). Some of the highlighted NN papers may refer to biological neural networks.

However, this simple analysis is constrained by the set of labels chosen a priori. The 2D map allows an unbiased exploration that does not rely on labels. For that, we highlighted all machine learning papers (*n* = 38, 446) in the embedding (Figure 4b). Papers claiming to use machine learning were grouped in the map into several clusters, covering topics ranging from computational biology to healthcare data management. These ML papers were more prevalent in the life science half of the map (left) and rather rare in the medical part (right). Within the medical part of the corpus, ML papers were concentrated in several regions, such as analysis of tumor imaging (radiology) or cancer biomarkers (oncology).

To further explore the ML-heavy regions, we selected and manually labeled 12 of them (Figure 4b) and computed the fraction of papers mentioning specific ML and statistical methods (Table S1). We found that the usage of ML techniques varied strongly across regions. Deep learning and convolutional networks were prominent in the image segmentation region (with applications e.g. in microscopy). Clustering was often used in analyzing sequencing data. Neural networks and support vector machines were actively used in structural biology. Principal component analysis was important for data analysis in mass spectrometry.

We expanded this analysis to the whole corpus by identifying 342,070 papers (1.7%) mentioning the same ML and statistical methods in their abstracts (Figure 4c). We found that the medical part of the embedding was dominated by classical linear methods such as linear regression and factor analysis, whereas more modern non-linear and nonparametric methods were mostly used in non-medical research. This shows that the medical disciplines are being slower in uptaking new computational techniques compared to basic life sciences.

### 2.5 Exploring the gender gap

In this section we will show how the map can be used to explore and better understand social disparities in biomedical publishing such as the extent and distribution of the well-known gender imbalance in academic authorship (Filardo et al., 2016; Larivière et al., 2013; Shen et al., 2018; Dworkin et al., 2020; Bendels et al., 2018). We used the first name (where available) of the first and the last author of every PubMed paper to infer their gender using the `gender` tool (Blevins and Mullen, 2015). The gender inference is only approximate, as many first 5 names were absent in the US-based training data, biasing our analysis towards Western academia, and some names are inherently gender-ambiguous (see Methods). Overall, this procedure allowed us to infer the gender of 62.3%/63.1% first/last authors with available first names. Among those, 42.4% of first authors and 29.1% of last authors were female. While some academic fields, such as mathematics and physics, tend to prefer alphabetic ordering of the authors, in biomedicine the first author is usually the trainee (PhD student or postdoc) who did the the practical hands-on project work and the last author is the supervisor or principal investigator.

The fraction of female authors steadily increased with time (Figure 5a), with first and last authors being 47.2% and 34.4% female in 2021. We found a delay of ∼20 years between the first and the last author curves, suggesting that it takes more than one academic generation for the differences in gender bias to propagate from mentees to mentors.

**Figure 5:**
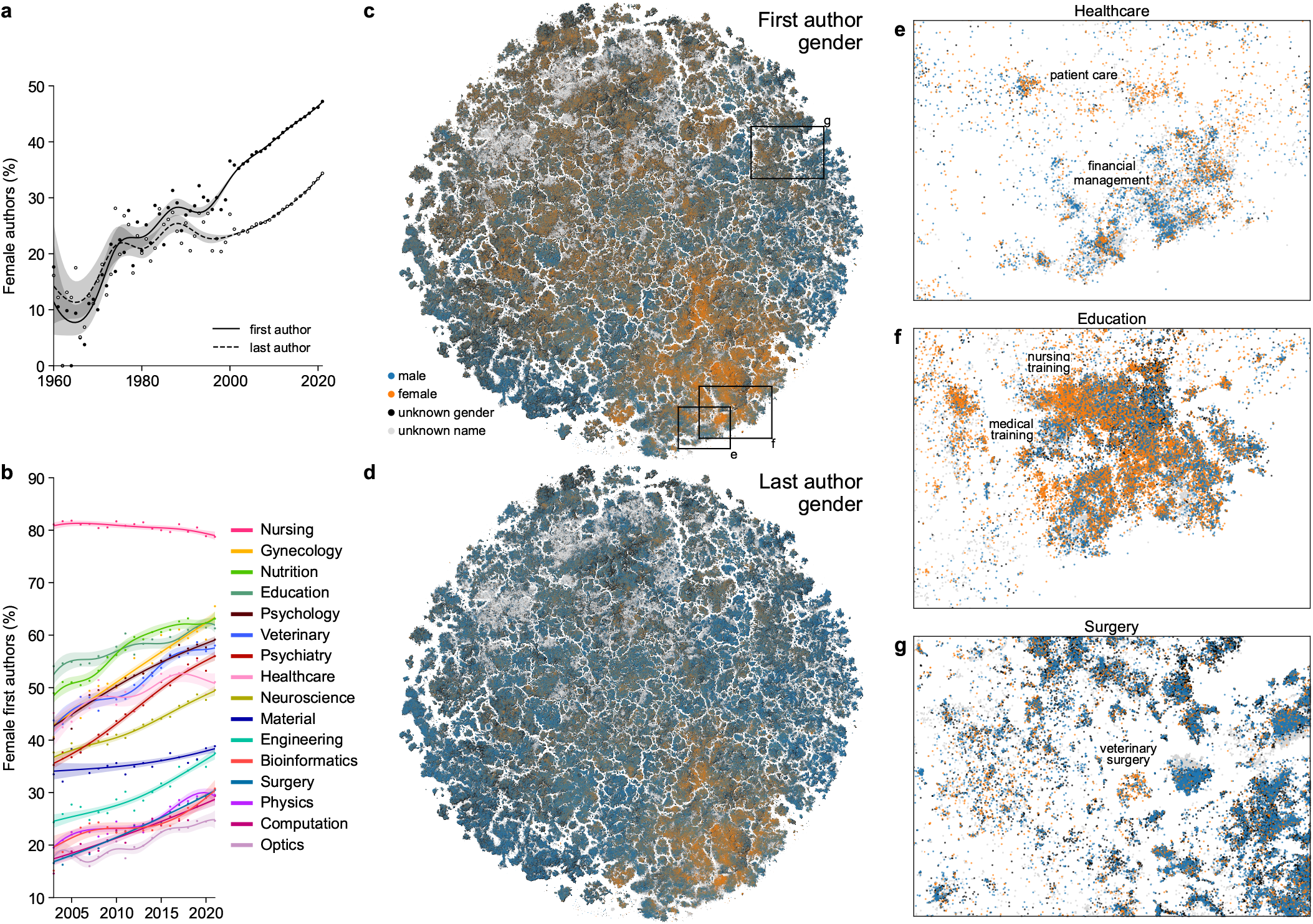
Gender bias in academic authorship. **(a)** Fraction of female first and last authors across time. The amount of available first names increased dramatically after 2003 (Figure S1c). Smooth curves and confidence intervals were obtained using generalized additive models (see Methods). **(b)** Fraction of female first authors across time for different disciplines. **(c)** Papers colored by the inferred gender of their first authors. **(d)** Papers colored by the inferred gender of their last authors. **(e–g)** Regions of the map showing within-label heterogeneity in the distribution of first authors’ gender: in healthcare (e), education (f), and surgery (g). Only papers belonging to those labels are shown.

Within most individual disciplines, the fraction of female first authors increased with time (Figure 5b), even in disciplines where this fraction was already high, such as education (increased from 55% female in 2005 to 60% in 2020). This increase also happened in male-dominated fields such as computation, physics, or surgery (increase from 15–20% to 25%). Notably, the female proportion in material sciences showed only a modest increase while nursing, the most female-dominated discipline across all our labels (80.4%) showed a moderate decrease.

Our map, when colored by gender, also showed that female authors were not equally distributed across the biomedical publishing landscape (Figure 5c). First and last female authors were most frequent in the lower right corner of the embedding, covering fields such as nursing, education, and psychology. Furthermore, the map allowed us to explore gender bias beyond the discipline level, revealing a substantial heterogeneity of gender ratios within individual disciplines. For example, in health-care (overall 49.6% female first authors), there were male- and female-dominated regions in the map. One of the more male-dominated clusters (33.9% female) focused on financial management while one of the more female ones (68.1% female) — on patient care (Figure 5c). In education (58.6% female authors), female authors dominated research on nursing training whereas male authors were more frequent in research on medical training (Figure 5d). In surgery, only 24.4% of the first authors were female, but this fraction increased to 61.1% in the cluster of papers on veterinary surgery (Figure 5e). This agrees with veterinary medicine being a predominantly female discipline (52.2% in total, Figure 5g). Importantly, these details are lost when averaging across a priori labels, while the embedding can suggest the relevant level of granularity.

### 2.6 Retracted papers highlight suspicious literature

We identified 11,756 papers flagged as retracted by PubMed and still having intact abstracts (not containing words like “retracted” or “withdrawn”; see Methods). These papers were not distributed uniformly over the 2D map (Figure 6) but instead concentrated in several specific areas, in particular on top of the map, covering research on cancer-related drugs, marker genes, and microRNA. These areas are known targets of paper mills (Byrne and Labbé, 2017; Byrne et al., 2019; Candal-Pedreira et al., 2022), which are organizations that produce fraudulent research papers for sale.

**Figure 6:**
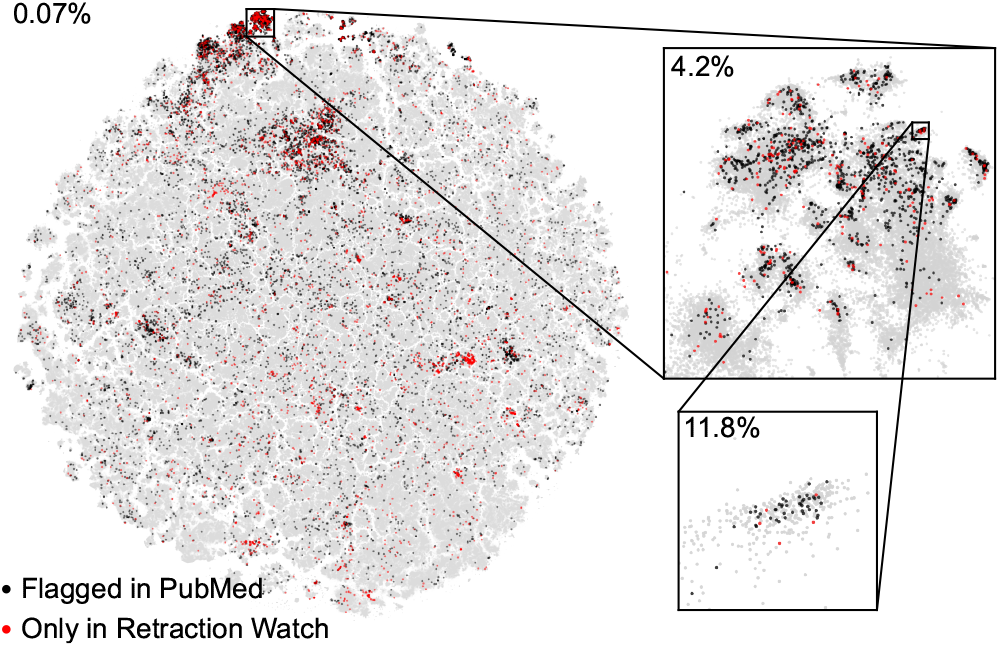
Retracted papers group together. All papers flagged as retracted by PubMed with intact abstracts (11,756) are highlighted in black, plotted on top of the non-retracted papers. Additional retracted papers (3,572) from the Retraction Watch database are shown in red. First inset corresponds to one of the regions with higher density of retracted papers (4.2%), covering research on cancer-related drugs, marker genes, and microRNA. Second inset corresponds to a subregion with a particularly high fraction of retracted papers (11.8%), the one we used for manual inspection.

Our map is based solely on textual similarity between abstracts. This suggests that non-retracted papers from the regions with high concentration of retracted papers may require an investigation, as their abstracts are similar to the ones from paper mill products. As an example, we considered a region with particularly high fraction (45/422) of retracted papers (second inset in Figure 6) and randomly selected 25 non-retracted papers for manual inspection. They had similar title format (variations of “MicroRNA-X does Y by targeting Z in osteosarcoma”; see Bielack and Palmerini, 2022), paper structure, and figure style, and 24/25 of them had authors affiliated with Chinese hospitals — features that are often shared by paper mill products (Byrne, 2019; Byrne and Christopher, 2020; Else and Van Noorden, 2021; Zhao et al., 2021; Candal-Pedreira et al., 2022; Fanelli et al., 2022; Sabel et al., 2023). Moreover, many areas with high fraction of retractions consisted of papers stemming mostly from a single country, typically China (Figure S8), which could by itself be an indicator of paper mill activity.

After we conducted our analysis, the Retraction Watch database of retracted papers was made open to the public. Using their database, we identified additional 3,572 papers in our map that were not marked as retracted in PubMed but were in fact retracted (red dots in Figure 6). They were mostly located in the same areas of the map that we identified as suspicious above, validating our conclusions. This does not guarantee that all papers in these areas are fraudulent, but confirms that our 2D map can be used to highlight papers requiring further editorial investigation (Oransky et al., 2021). If additional paper mills are discovered in the future, our map will help to highlight literature clusters requiring further scrutiny.

## 3 Discussion

We developed a two-dimensional atlas of the biomedical literature based on the PubMed collection of 21 M paper abstracts using a transformer-based language model (PubMedBERT) and a neighbor embedding visualization (*t*-SNE) tailored to handle large document libraries. We used this atlas as an exploration tool to study the biomedical research landscape, generating hypotheses that we later confirmed using the original high-dimensional data. Using five distinct examples — the emergence of the Covid-19 literature, the evolution of the neuroscience discipline, the uptake of machine learning, the gender imbalance, and the concentration of retracted fraudulent papers — we argued that two-dimensional visualizations of text corpora can help uncover aspects of the data that other analysis methods may fail to reveal.

We also developed an interactive web version of the embedding (https://static.nomic.ai/pubmed.html) based on the deepscatter library (Nomic AI, 2022) that allows to navigate the atlas, zoom, and search by title, journal, or author names. In deepscatter, individual points are loaded on demand when zooming-in, like when navigating geographical maps in the browser. This interactive website contains a separate embedding of the latest PubMed data, including 2022–2023 papers (Figures S9– S10). We plan on updating the visualization in the future using annual PubMed releases.

Neighbor embedding methods like *t*-SNE have known limitations. For the datasets of our size, the few closest neighbors in the two-dimensional embedding space are typically different from the neighbors in the high-dimensional BERT representation (Table 1). This makes our map suboptimal for finding the most similar papers to a given query paper, and other tools, like conventional (Google Scholar, PubMed) or citation-based (`connectedpapers.com`) search engines, may be more appropriate for this task. Instead, our map is useful for navigating the literature on the scale of narrow and focused scientific topics. Neighbor embedding algorithms can misrepresent the global organization of the data (Wattenberg et al., 2016; Kobak and Berens, 2019; Böhm et al., 2022). We used methods designed to mitigate this issue (Kobak and Berens, 2019; Kobak and Linderman, 2021; González-Márquez et al., 2022), and indeed, found that related research areas were located close to each other.

In annotating our atlas, we selected 38 labels spanning various life science fields and medical specialties. Each label was assigned to papers in journals with the corresponding word in their titles, resulting in 34.4% of papers being labeled. Although this method overlooks interdisciplinary journals like *Science* or *Nature*, it ensures that core works in each discipline are labeled. While PubMed uses several systems to organize articles into categories (such as keywords or Medical Subject Headlines [MeSH] terms), creating labels based on them would likely require a more involved manual curation process. We found that our labels covered most of the two-dimensional space (Figure 1), and therefore considered the labeled subset representative of the entire landscape.

Our atlas provides the most detailed visualization of the biomedical literature landscape to date. Previously, PubMed abstracts were clustered based on textual bag-of-words similarity and citation information, and the clusters were displayed using a two-dimensional embedding (Boyack et al., 2020). Their map exhibits similar large-scale organization, but only shows 29,000 clusters, so our map is almost three orders of magnitude more detailed. The BioBERT model was previously applied to the PubMed dataset to extract information on biomedical concepts, such as proteins or drugs (Xu et al., 2020). Previous work on visualizing large text corpora includes Schmidt (2018) and González-Márquez et al. (2022). Both were based on bag-of-words representations of the data. Here we showed that BERT-based models outperform TF-IDF for representing scientific abstracts.

An alternative approach to visualizing collections of academic works is to use information on citations as a measure of similarity, as opposed to semantic or textual similarity. For example, `paperscape.org` visualizes 2.2 M papers from the *arXiv* preprint server using a force-directed layout of the citation graph. Similarly, `opensyllabus.org` uses `node2vec` (Grover and Leskovec, 2016) and UMAP to visualize 1.1 M texts based on their co-appearance in the US college syllabi. Similar approach was used by Noichl (2021) to visualize 68,000 articles on philosophy based on their reference lists. Here we based our embedding on the abstract texts alone, and in future work it would be interesting to combine textual and co-citation similarity in one map (citation graph for PubMed papers can be obtained from OpenAlex (Priem et al., 2022), MAG (Sinha et al., 2015), and/or PubMed itself). The functionality of our interactive web version is similar to `opensyllabus.org` and `paperscape.org`, but we successfully display one order of magnitude more points.

We achieved the best representation of the PubMed abstracts using the PubMedBERT model. As the progress in the field of language models is currently very fast, it is likely that a better representation may soon become available. One promising approach could be to train sentence-level models such as SBERT (Reimers and Gurevych, 2019) on the biomedical text corpus. Another active avenue of research is fine-tuning BERT models using contrastive learning (Gao et al., 2021; Liu et al., 2021) and/or using citation graphs (Cohan et al., 2020; Ostendorff et al., 2022). While we found that these models were outperformed by PubMedBERT, similar methods (Yasunaga et al., 2022) could be used to fine-tune the PubMedBERT model itself, potentially improving its representation quality further. Finally, larger generative language models such as recently developed BioGPT (Luo et al., 2022) or BioMedLM (Stanford CRFM and MosaicML, 2022) can possibly lead to better representations as well.

In conclusion, we suggested a novel approach for visualizing large document libraries and demonstrated that it can facilitate data exploration and help generate novel insights. Many further meta-scientific questions can be investigated in the future using our approach.

## 4 Methods

### 4.1 PubMed dataset

We downloaded the complete PubMed database (295 GB) as XML files using the bulk download service (www.nlm.nih.gov/databases/download/pubmed_medline.html). PubMed releases a new snapshot of their database every year; they call it a ‘baseline’. In our previous work (González-Márquez et al., 2022) we used the 2020 baseline (files called `pubmed21n0001.xml.gz` to `1062.xml.gz`, download date: 26.01.2021). In this work, we supplemented them with the additional files from the 2021 baseline (files called `pubmed22n1062.xml.gz` to `1114.xml.gz`, download date: 27.04.2022). After the analysis was completed, we realized that our dataset had 0.07% duplicate papers; they should not have had any noticeable influence on the reported results.

We used the Python `xml` package to extract PubMed ID, title, abstract, language, journal title, ISSN, publication date, and author names of all 33.4 M papers. We filtered out all 4.7 M non-English papers, 10.8 M papers with empty abstracts, 0.3 M papers with abstracts shorter than 250 or longer than 4000 symbols (Figures S1, S11), and 27 thousand papers with unfinished abstracts. Papers with unfinished abstracts needed to be excluded because otherwise they were grouped together in the BERT representation, creating artifact clusters in the embedding. We defined unfinished abstracts as abstracts not ending with a period, a question mark, or an exclamation mark. Some abstracts ended with a phrase “(ABSTRACT TRUNCATED AT … WORDS)” with a specific number instead of ‘…’. We removed all such phrases and analyzed the remaining abstracts as usual, even though they did not contain the entire text of the original abstracts. In some cases, abstracts were divided in subsections (such as Methods, Results, etc.). We excluded subsection titles so that the resulting abstract had effectively a single paragraph. Overall, we were left with 20,687,150 papers for further analysis.

This collection contains papers from the years 1808– 2022. MEDLINE, the largest component of PubMed, started its record in 1966 and later included some noteworthy earlier papers. Therefore, the majority (99.8%) of the PubMed papers are post-1970 (Figure S1c). There are only few papers from 2022 in our dataset. The 2021 data in this PubMed snapshot were also incomplete.

### 4.2 Label assignment

We labeled the dataset by selecting 38 keywords contained in journal titles that reflected the general topic of the paper. We based our choice of keywords on lists of medical specialties and life science branches that appeared frequently in the journal titles in our dataset. The 38 terms are: anesthesiology, biochemistry, bioinformatics, cancer, cardiology, chemistry, computation, dermatology, ecology, education, engineering, environment, ethics, genetics, gynecology, healthcare, immunology, infectious, material, microbiology, neurology, neuroscience, nursing, nutrition, ophthalmology, optics, pathology, pediatric, pharmacology, physics, physiology, psychiatry, psychology, radiology, rehabilitation, surgery, veterinary, and virology.

Papers were assigned a label if their journal title contained that term, either capitalized or not, and were left unlabeled otherwise. Journal titles containing more than one term were assigned randomly to one of them. This resulted in 7,123,706 labeled papers (34.4%).

Our journal-based labels do not constitute the ground truth for the topic of each paper, and so the highest possible classification accuracy is likely well below 100%. Nevertheless, we reasoned that the higher the classification accuracy, the better the embedding, and found this metric to be useful to compare different representations (Tables 1, 3).

### 4.3 BERT-based models

We used PubMedBERT (Gu et al., 2021) to obtain a numerical representation of each abstract. Specifically, we used the HuggingFace’s `transformers` library and the publicly released PubMedBERT model. PubMedBERT is a Transformer-based language model trained in 2020 on PubMed abstracts and full-text articles from PubMed Central.

In pilot experiments, we compared performance of eight BERT variants: the original BERT (Devlin et al., 2019), SciBERT (Beltagy et al., 2019), BioBERT (Lee et al., 2020), PubMedBERT (Gu et al., 2021), SBERT (Reimers and Gurevych, 2019), SPECTER (Cohan et al., 2020), SimCSE (Gao et al., 2021), and SciNCL (Ostendorff et al., 2022). The exact HuggingFace models that we used:

- `bert-base-uncased`
- `allenai/scibert scivocab uncased`
- `dmis-lab/biobert-v1.1`
- `microsoft/BiomedNLP-PubMedBERT-base-uncased-abstract-fulltext`
- `sentence-transformers/all-mpnet-base-v2`
- `allenai/specter`
- `malteos/scincl`
- `princeton-nlp/unsup-simcse-bert-base-uncased`

All of these models have the same architecture (`bert-base`; 110M parameters) but were trained and/or fine-tuned on different data. The original BERT was trained on a corpus of books and text from Wikipedia. SciBERT was trained on a corpus of scientific articles from different disciplines. BioBERT fine-tuned the original BERT on PubMed abstracts and full-text articles from PubMed-Central. PubMedBERT was trained on the same data from scratch (and its vocabulary was constructed from PubMed data, whereas BioBERT used BERT’s vocabulary).

The other four models were fine-tuned to produce sentence embeddings instead of word embeddings, i.e. to generate a single vector representation of the entire input text (we treated each entire abstract as one single ‘sentence’ when providing it to these models). SBERT fine-tuned BERT using a corpus of similar sentences and paragraphs; the specific model that we used was obtained via fine-tuning MPNet (Song et al., 2020). According to SBERT’s authors, this is currently the most powerful generic SBERT model; note that their training procedure has evolved since the original approach described in Reimers and Gurevych (2019). SPECTER and SciNCL, both fine-tuned the SciBERT model using contrastive loss functions based on the citation graph. SimCSE fine-tuned the original BERT using a contrastive loss function between the sentence representations obtained with two different dropout patterns, using Wikipedia texts.

For this pilot experiment, we used a subset of our data (*n* = 1,000,000 labeled papers; 990,000 were used as a training set and 10,000 as a test set) to measure *k*NN accuracy (*k* = 10) of each of these models, and obtained the highest accuracy with PubMedBERT (see Table 3). This made sense as PubMedBERT’s training data largely overlapped with our dataset. We found that SBERT performed better than BERT, but did not reach the level of PubMedBERT on our task. SimCSE did not outperform the original BERT in our benchmark. SPECTER and SciNCL outperformed SciBERT, suggesting that citation information can be helpful for training scientific language models. Still, both models performed worse than PubMedBERT on our task.

Furthermore, we compared *k*NN accuracy after *t*-SNE between different BERT models (Figure S12), and again obtained the best results with PubMedBERT (Table 4). The performance of SciNCL here was only 0.1% lower. We used the same settings for *t*-SNE as described below, but ran it with the default number of iterations (750).

**Table 4:** *k*NN accuracy of *t*-SNE representations of different BERT-based models. The same experimental setup as in Table 3. For comparison, the accuracy of *t*-SNE of the TF-IDF representation (after SVD) was 49.9%.

|  | Average | [CLS] | [SEP] |
| --- | --- | --- | --- |
| BERT | 46.0% | 36.3% | 40.6% |
| SciBERT | 52.3% | 43.4% | 48.8% |
| BioBERT | 54.7% | 51.1% | 56.5% |
| PubMedBERT | 53.2% | 45.5% | **60.8%** |
| SBERT | 60.2% | 56.3% | 56.7% |
| SPECTER | 58.4% | 59.2% | 59.3% |
| SciNCL | 60.7% | 59.1% | 59.4% |
| SimCSE | 46.9% | 42.4% | 40.3% |

Each abstract gets split into a sequence of tokens, and PubMedBERT represents each token in a 768-dimensional latent space. PubMedBERT’s maximum input length is 512 tokens and longer abstracts are automatically truncated at 512 tokens (this corresponds to roughly 300–400 words, and ∼98% of all abstracts were shorter than 512 tokens). We are interested in a single 768-dimensional representation of each abstract, rather than 512 of them. For this, we compared several approaches commonly used in the literature: using the representation of the initial `[CLS]` token, the trailing `[SEP]` token, and averaging the representations of all tokens (Devlin et al., 2019; Reimers and Gurevych, 2019; Beltagy et al., 2019). Using the `[SEP]` token yielded the highest *k*NN accuracy in our pilot experiments (Table 3), so we adopted this approach.

Note that sentence transformers were originally trained to optimize one specific representation, e.g. SBERT uses the average representation across all tokens as its sentence-level output, while SPECTER uses the `[CLS]` token. For consistency, in Table 3 we report the performance of all three representations for each model. SBERT implementation (`sentence-transformers` library) normalizes its output to have norm 1. In Table 3 we report the accuracy without this normalization (64.5%), as obtained using the `transformers` library; with normalization, the accuracy changed by less than 0.1%.

Su et al. (2021) argued that whitening BERT representation can lead to a strongly improved performance on some benchmarks. We tried whitening the PubMed-BERT representation, but only observed a decrease in the *k*NN accuracy. For this experiment, we used a test set of 500 labeled papers, and compared PubMedBERT without any transformations, after centering, and after whitening, using both Euclidean metric and the cosine metric, following Su et al. (2021). We obtained the best results using the raw PubMedBERT representation (Table 5). Our conclusion is that whitening does not improve the *k*NN graph of the PubMedBERT representation.

**Table 5:** *k*NN accuracy of label prediction using different transformations of the PubMedBERT representation and two different metrics for finding nearest neighbors. This experiment used test set size 500, smaller than in Table 1.

|  | Euclidean | Cosine |
| --- | --- | --- |
| **Raw** | **67.8%** | **67.8%** |
| Centered | 67.8% | 67.4% |
| Whitened | 64.2% | 65.4% |

In the end, our entire collection of abstracts is represented as a 20,687,150 *×* 768 dense matrix.

### 4.4 TF-IDF representation

In our prior work (González-Márquez et al., 2022), we used the bag-of-words representation of PubMed abstracts and compared several different normalization approaches. We obtained the highest *k*NN accuracy using the TF-IDF (term frequency inverse document frequency) representation (Jones, 1972) with log-scaling, as defined in the scikit-learn implementation (version 0.24.1):

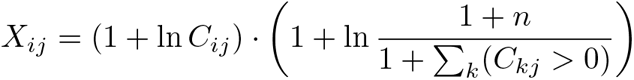

if *C*_*ij*_ *>* 0 and *X*_*ij*_ = 0 otherwise. Here *n* is the total number of abstracts and *C*_*ij*_ are word counts, i.e. the number of times word *j* occurs in abstract *i*. In the scikit-learn implementation, the resulting *X*_*ij*_ matrix is then row-normalized, so that each row has 𝓁_2_ norm equal to 1.

This results in a 20,687,150 × 4,679,130 sparse matrix with 0.0023% non-zero elements, where 4,679,130 is the total number of unique words in all abstracts.

This matrix is too large to use in *t*-SNE directly, so for computational convenience we used truncated SVD (`sklearn.decomposition.TruncatedSVD` with `algorithm=‘arpack’`) to reduce dimensionality to 300, the largest dimensionality we could obtain given our RAM resources. Note that we did not use SVD when using BERT representations and worked directly with 768-dimensional representations.

The *k*NN accuracy values for the TF-IDF and SVD (*d* = 300) representations measured on the same 1 M subset as used in the previous section were 61.0% and 54.8% respectively. After *t*-SNE, the *k*NN accuracy were 49.9%. We also tried row-normalizing the SVD representation and observed that this increased the kNN accuracy to 58.7% (and 52.0% after *t*-SNE); this is equivalent to using cosine distance instead of Euclidean distance for finding nearest neighbors.

We have also experimented with constructing a TF-IDF representation based on the PubMedBERT’s tokenizer, instead of the default TF-IDF tokenizer. This reduces the vocabulary size and the dimensionality of the resulting space from 758,111 to 29,047 (because PubMed-BERT’s tokenizer does not include all unique words as tokens, but instead fragments rare words into repeating sub-strings). This barely affected *k*NN classification accuracy: it changed from 61.0% to 61.6% in the high-dimensional space and from 49.9% to 50.1% in the two-dimensional space after SVD and *t*-SNE. Note that the actual PubMedBERT representation captures many more aspects of the text than just the presence or absence of specific tokens, so it is unsurprising that the representation quality was higher there (67.7% in 768D and 60.8% in 2D).

### 4.5 *t*-SNE

We used the `openTSNE` (version 0.6.0) implementation (Poličar et al., 2019) of *t*-SNE (van der Maaten and Hinton, 2008) to reduce dimensionality from 768 (for the BERT representation) or 300 (for the TF-IDF representation) to *d* = 2. OpenTSNE is a Python reimplementation of the FIt-SNE (Linderman et al., 2019) algorithm.

We ran *t*-SNE following the procedure established in our prior work (González-Márquez et al., 2022): using uniform affinities (on the approximate *k*NN graph with *k* = 10) instead of perplexity-based affinities, early exaggeration annealing instead of the abrupt switch of the early exaggeration value, and extended optimization for 2250 iterations instead of the default 750 (250 iterations for the early exaggeration annealing, followed by 2000 iterations without exaggeration). We did not use any ‘late’ exaggeration after the early exaggeration phase. All other parameters were kept at default values, including PCA initialization and learning rate set to *n/*12, where *n* is the sample size.

In our previous work we showed that this visualization approach outperformed UMAP (version 0.5.1) (McInnes et al., 2018) on PubMed data in TF-IDF representation in terms of both *k*NN recall and *k*NN accuracy (González-Márquez et al., 2022). We confirmed that the same was true for the PubMedBERT representation of the 1 M subset used in the previous sections (Figure S13): the UMAP embedding was qualitatively similar to the *t*-SNE embedding with exaggeration *ρ* = 4, and its *k*NN recall (2.8%) and accuracy (49.6%) were lower than those obtained using *t*-SNE without exaggeration (12.5% and 60.8% respectively).

The *t*-SNE embeddings of a PubMed subset containing 1 million papers (Figures S12, S13, Table 4) used the default number of iterations (750).

The embeddings based on the TF-IDF and PubMed-BERT representation showed similar large-scale organization. As *t*-SNE loss function is unaffected by rotations and/or sign flips, we flipped the *x* and/or *y* coordinates of the TF-IDF *t*-SNE embedding to match its orientation to the PubMedBERT *t*-SNE embedding. The same was done for the embeddings shown in Figures S12, S13.

### 4.6 Performance metrics

All *k*NN-based metrics were based on *k* = 10 exact nearest neighbors, obtained using the `NearestNeighbors` and `KNeighborsClassifier` classes from scikit-learn (version 1.0.2) using `algorithm=‘brute’` and `n_jobs=-1` (Pedregosa et al., 2011).

To predict each test paper’s label, *k*NN classifier takes the majority label among the paper’s nearest neighbors in the training set. To measure the accuracy, the classifier was trained on all labeled papers excluding a random test set of labeled papers. The test set size was 5000 for the high-dimensional representations and 10000 for the two-dimensional ones. The chance-level *k*NN accuracy was obtained using the `DummyClassifier` from scikit-learn with `strategy=‘stratified’`, and test set size 10000.

To predict each test paper’s publication year, we took the average publication year of the paper’s nearest neighbors in the training set. To measure the root-mean-squared error (RMSE), we used the training set consisting of all papers excluding a random test set. The test set size was 5000 for the high-dimensional representations and 10000 for the two-dimensional ones. The chance-level root-mean-squared error (RMSE) was calculated by drawing 10 random papers instead of nearest neighbors, for a test set of 5000 papers.

We define *k*NN recall as the average size of the overlap between *k* nearest neighbors in the high-dimensional space and *k* nearest neighbors in the low-dimensional space. We averaged the size of the overlap across a random set of 10000 papers for the BERT representation, and 5000 papers for the TF-IDF representation. The *k*NN recall value reported in Table 1 for the TF-IDF representation measures the recall of the original TF-IDF neighbors (0.7%); the recall of the neighbors from the SVD space (which was used for *t*-SNE) was 1.5%.

Isolatedness metric was defined as the average fraction of *k* nearest neighbors belonging to the same corpus. We used a random subset of 5000 papers from each corpus to estimate the isolatedness. The regions from Table 2 were selected as follows. The HIV/AIDS set contained all papers with ‘HIV’ or ‘AIDS’ words (upper case or lower case) appearing in the abstract. The influenza set contained all papers with the word ‘influenza’ in the abstract (capitalized or not). Similarly, meta-analysis set was obtained using the word ‘meta-analysis’. The virology and ophthalmology sets correspond to the journal-based labels (see above).

### 4.7 Covid-related papers

We considered a paper Covid-related if it contained at least one of the following terms in its abstract: ‘covid-19’, ‘COVID-19’, ‘Covid-19’, ‘CoViD-19’, ‘2019-nCoV’, ‘SARS-CoV-2’, ‘coronavirus disease 2019’, ‘Coronavirus disease 2019’. Our dataset included 132,802 Covid-related papers.

We selected 27 frequent terms contained in Covid-related paper titles to highlight different subregions of the Covid cluster. The terms were: antibody, anxiety, cancer, children, clinical, epidemic, healthcare, immune, implications, mental, mortality, outbreak, pediatric, pneumonia, population, psychological, respiratory, social, strategies, students, surgery, symptoms, therapy, transmission, treatment, vaccine, and workers. Papers were assigned a keyword if their title contained that term, either capitalized or not. Paper titles containing more than one term were assigned randomly to one of them. This resulted in 35,874 Covid-related papers containing one of those keywords: 27.0% from the total amount of Covid-related papers and 45.6% of the Covid-related papers from the main Covid cluster in the embedding.

### 4.8 Generalized additive models

We used generalized additive models (GAMs) to obtain smooth trends for several of our analyses across time (Figures 3c, 4c, 5c–d). We used the `LinearGAM` (GAM with the Gaussian error distribution and the identity link function) and the `LogisticGAM` (GAM with the binomial error distribution and the logit link function) from the `pyGAM` Python library (version 0.8.0) (Servén and Brummitt, 2018). In all cases, we excluded papers published in 2022, since we only had very few of them (as we used the 2021 baseline of the PubMed dataset, see above). Linear GAMs (with `n_splines=6`) were used for Figure 3c, and logistic GAMs (with `n_splines=12`) were used for Figures 4c and 5c–d. All GAMs had the publication year as the only predictor.

In all cases, we used the `gridsearch()` function to estimate the optimal smoothing (lambda) parameter using cross-validation. To obtain the smooth curves shown in the plots, we predicted the dependent value on a grid of publication years. The confidence intervals were obtained using the `confidenceintervals()` function from the same package.

In Figure 3c, the response variable was *k*NN overlap of a neuroscience paper with the target discipline. For each discipline, the input data was a set of 500 randomly chosen neuroscience papers for each year in 1975–2021. If the total number of neuroscience papers for a given year was less than 500, all of them were taken for the analysis. The *k*NN overlap values of individual papers were calculated using *k* = 10 nearest neighbors obtained with the `NearestNeighbors` class.

In Figure 4c, the binary response variable was whether a paper contained ‘machine learning’ in its abstract. For each discipline, the input data were all 2010–2021 papers. In Figure 5c–d, the binary response variable was whether the paper’s first or last author was female (as inferred by the `gender` tool, see below). The input data in all cases were all papers with gender information from 1960–2021.

### 4.9 Gender prediction

We extracted authors’ first names from the XML tag `ForeName` that should in principle only contain the first name. However, we observed that sometimes it contained the full name. For that reason, we always took the first word of the `ForeName` tag contents (after replacing hyphens with spaces) as the author’s first name. This reduced some combined first names (such as Eva-Maria or Jose Maria) to their initial word (Eva; Jose). In many cases, mostly in older papers, the only available information about the first name was an initial. As it is not possible to infer gender from an initial, we discarded all extracted first names with length 1. In the end we obtained 13,429,169 first names of first authors (64.9% of all papers) and 13,189,271 first names of last authors (63.8%), almost only from 1960–2022.

We used the R package `gender` (Blevins and Mullen, 2015) (version 0.6.0) to infer authors’ genders. This package uses a historical approach that takes into account how naming practices have changed over time, e.g., Leslie used to be a male name in the early XX century but later has been mainly used as a female name. For each first/last author, we provided `gender` with the name and the publication year, and obtained the inferred gender together with a confidence measure.

The `gender` package offers predictions based on different training databases. We used the 1930–2012 Social Security Administration data from United States (`method=‘ssa’`). For the papers published before 1930 we fixed the year to 1930 and for the papers published after 2012, we fixed it to 2012. The SSA data do not contain information on names that are not common in the USA, and we only obtained inferred genders for 8,363,116 first authors (62.3% of available first names) and 8,468,165 last authors (63.1% of available last names). Out of all inferred genders, 3,543,592 first authors (42.4%) and 2,464,882 last authors (29.1%) were female.

Importantly, our gender inference is only approximate (Blevins and Mullen, 2015). The inference model has clear limitations, including limited US-based training data and state-imposed binary genders. Moreover, some first names are inherently gender-ambiguous. However, the distribution of inferred genders over biomedical fields and the pattern of changes over the last decades matched what is known about the gender imbalance in academia, suggesting that inferred genders were sufficiently accurate for our purposes.

### 4.10 Retracted papers

We obtained PMIDs of papers classified in PubMed as retracted (13,569) using the PubMed web interface on 19.04.2023. Of those, 11,998 were present in our map (the rest were either filtered out in our pipeline or not included in the 2021 baseline dataset we used). To make sure that retracted papers were not grouping together in the BERT space because their abstract had been modified to indicate a retraction, we excluded from consideration all retracted papers containing the words “retracted”, “retraction”, “withdrawn”, or “withdrawal” in their abstract (242 papers). The remaining retracted papers (11,756) had intact original abstracts and are shown in Figure 6.

There was one small island at the bottom of the map containing retraction notices (they have independent PubMed entries with separate PMIDs) as well as corrigenda and errata, which were not filtered out by our length cutoffs. Many of the 242 retracted papers with post-retraction modified abstracts were also located there. We obtained the Retraction Watch database through (https://api.labs.crossref.org/data/retractionwatch?name@email.org) as a CSV file (41 MB) on 21.09.2023. It contained 18,786 retracted papers indexed in PubMed. Of those, 15,666 were present in our map (the rest were either filtered out in our pipeline or not included in the 2021 baseline dataset we used). 15,103 of those were intact papers. These 15,103 papers contained all of the 11,998 papers used above except for 234 papers. This gave 3,572 additional retracted papers shown in Figure 6 in red.

### 4.11 2023 annual PubMed baseline

While the paper was in revision, we updated the dataset by downloading the latest annual PubMed snapshot (2023 baseline; files called `pubmed24n0001.xml.gz` to `1219.xml.gz`, download date: 06.02.2024, 350 GB). We used this entire dataset, and not only the files containing 2022–2023 papers, to avoid duplicated entries and to use the latest meta-data. This snapshot included in total 36,555,430 papers. After filtering with our previous criteria, we were left with 23,389,083 papers.

We extracted the same attributes from the metadata as described above, with the addition of the first affiliation of the first author. We used this affiliation to assign each paper to a country (Figure S10), by searching the string for all existing country names in English (taking into account possible name variations, such as “United Kingdom” and “UK”). Consequently, papers that may have included country names in their original language (e.g. “Deutschland” instead of “Germany”) were not matched to any country. We noticed that many US affiliations did not explicitly include country name so we assigned affiliations containing a name of any US state to the US. This resulted in 19,937,913 papers (85.2%) with an assigned country. We matched the affiliation countries to our main dataset (2021 baseline) using PMIDs, which led to 17,404,977 papers (84.1%) with an assigned country (Figure S8).

We used the same journal-based labels to color the embedding (Figure S9) and added ‘dentistry’ as an additional label. This resulted in 8,028,583 labeled papers (34.3%).

### 4.12 Runtimes

Computations were performed on a machine with 384 GB of RAM and Intel Xeon Gold 6226R processor (16 multi-threaded 2.90 GHz CPU cores) and on a machine with 512 GB of RAM and Intel Xeon E5-2630 v4 processor (10 multi-threaded 2.20 GHz CPU cores). BERT embeddings were calculated using an NVIDIA TITAN Xp GPU with 12.8 GB of RAM.

Parsing the XML files took 10 hours, computing the PubMedBERT embeddings took 74 hours, running *t*-SNE took 8 hours. More details are given in Table S2. We used exact nearest neighbors for all *k*NN-based quality metrics, so evaluation of the metrics took longer than computing the embedding. In total, it took around 8 days to compute all the reported metrics (Table S2).

## Data and code availability

The analysis code is available at https://github.com/berenslab/pubmed-landscape. All original code has been deposited at Zenodo under the DOI 10.5281/zen-odo.10727578 https://zenodo.org/records/10727578 and is publicly available as of the date of publication.

We made publicly available a processed version of our dataset: a `csv.zip` file (20,687,150 papers, 1.3 GB) including PMID, title, journal name, publication year, embedding *x* and *y* coordinates, our label, and our color used in Figure 1a. We also included two additional files: the raw abstracts (`csv.zip` file, 9.5 GB), and the 768-dimensional PubMedBERT embeddings of the abstracts (NumPy array in float16 precision, 31.8 GB). They can all be downloaded from https://zenodo.org/record/7695389.

## Acknowledgments

We thank Richard Van Noorden, David Bimler, Ivan Oransky, and Jennifer Byrne for discussions. This research was funded by the Deutsche Forschungsgemeinschaft (KO6282/2-1, BE5601/8-1, and EXC 2064 “Machine Learning: New Perspectives for Science”, 390727645), by the German Ministry of Education and Research (Tübingen AI Center), and by the Hertie Foundation. The authors thank the International Max Planck Research School for Intelligent Systems (IMPRS-IS) for supporting Rita González Márquez.

### A Appendix

#### A.1 Supplementary Tables

**Table S1:** Percentage of abstracts mentioning various machine learning methods (as in Figure 4a) in each region of the embedding with high fraction of abstracts mentioning ‘machine learning’ (Figure 4b). Percentages above 4% in bold. Rows ordered by the number of papers in the region. Columns ordered by the average percentage across regions. Abbreviations as in Figure 4.

| # | Region | NN | Clustering | DL | SVM | CNN | PCA | RF | LR | DR | FA |
| --- | --- | --- | --- | --- | --- | --- | --- | --- | --- | --- | --- |
| 1 | EEG signals | **6.1** | 1.7 | 1.9 | 3.3 | 2.1 | 1.1 | 0.8 | 0.8 | 0.4 | 0.2 |
| 2 | Sequencing | 1.5 | **5.8** | 1.0 | 0.9 | 0.7 | 0.4 | 0.6 | 0.3 | 0.3 | 0.1 |
| 3 | Image segmentation | **9.5** | 2.4 | **7.3** | 2.5 | **7.6** | 0.9 | 1.0 | 0.5 | 0.2 | 0.1 |
| 4 | ML algorithms | **14.7** | **5.7** | **4.3** | 2.9 | **5.1** | 1.1 | 0.6 | 0.5 | 0.9 | 0.2 |
| 5 | Mass spectrometry | 1.8 | 1.2 | 0.2 | 1.5 | 0.2 | **4.9** | 0.4 | 1.1 | 0.2 | 0.5 |
| 6 | Healthcare data | 1.7 | 0.8 | 1.6 | 0.6 | 0.6 | 0.0 | 0.2 | 0.0 | 0.0 | 0.0 |
| 7 | Cancer biomarkers | 0.5 | **4.9** | 0.2 | 1.0 | 0.1 | 1.1 | 0.8 | 0.3 | 0.1 | 0.1 |
| 8 | Protein structure | **5.5** | 3.5 | 1.9 | **4.6** | 1.0 | 0.7 | 1.7 | 1.4 | 0.3 | 0.1 |
| 9 | Computational chemistry | 1.9 | 0.4 | 0.3 | 0.1 | 0.2 | 0.1 | 0.1 | 0.2 | 0.1 | 0.0 |
| 10 | Tumor imaging | 3.3 | 1.2 | 2.8 | 3.6 | 2.0 | 0.7 | 2.6 | 1.3 | 0.3 | 0.1 |
| 11 | Microbiome | 0.1 | 2.9 | 0.0 | 0.2 | 0.0 | 1.2 | 1.5 | 0.9 | 0.0 | 0.1 |
| 12 | Covid-19 tweets | 1.5 | 2.1 | 1.9 | 0.7 | 0.4 | 0.2 | 0.8 | 0.6 | 0.0 | 0.0 |

**Table S2:** Runtimes for different analyses.

| Step | Time |
| --- | --- |
| Parsing XML | 10 h |
| PubMedBERT representation | 74 h |
| TF-IDF representation | 1 h |
| Truncated SVD of TF-IDF | 4 h |
| *t*-SNE affinities for PubMedBERT | 101 min |
| *t*-SNE affinities for TF-IDF | 78 min |
| *t*-SNE optimization, 750 iter. | 126 min |
| *t*-SNE optimization, 2250 iter. | 390 min |
| *k*NNs for 1k papers, BERT | 32 min |
| *k*NNs for 1k papers, BERT labeled | 7 min |
| *k*NNs for 1k papers, TF-IDF | 150 min |
| *k*NNs for 1k papers, TF-IDF labeled | 12 min |
| *k*NNs for 1k papers, *t*-SNE | 7 min |
| *k*NNs for 1k papers, *t*-SNE labeled | 2 min |
| Table 1 | ~35 h |
| Table 2 | ~91 h |
| Figure 3c | ~20 h |
| Table 3 (BERT computations) | ~30 h |
| Table 4 | ~7 h |
| Table 5 | ~20 min |
| All GAMs | ~30 min |
| Gender prediction | 6 min |

#### A.2 Supplementary Figures

**Figure S1:**
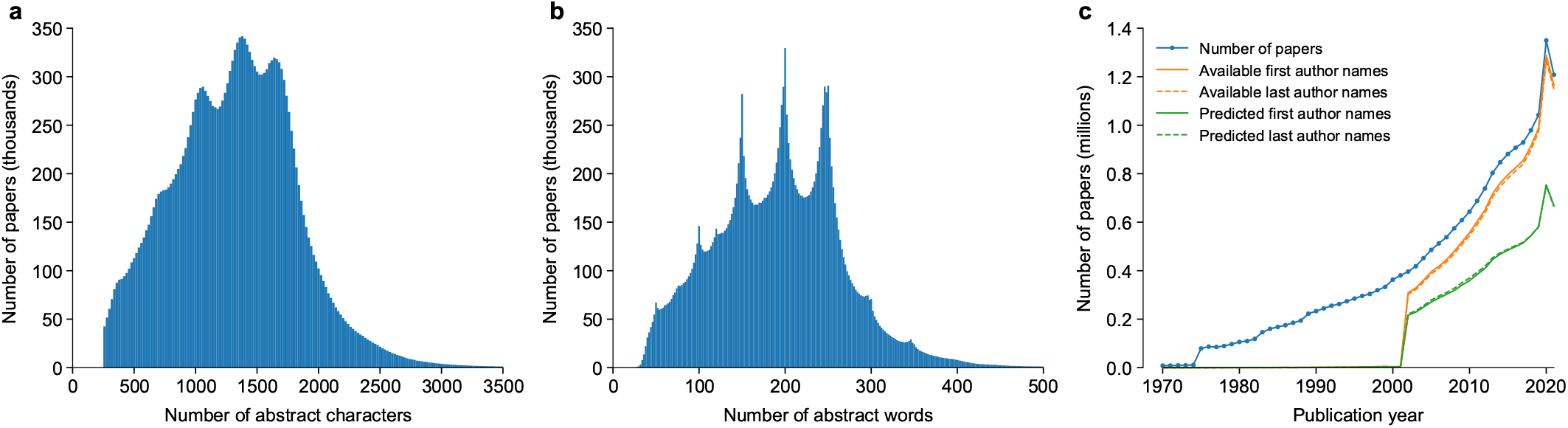
Summary of the PubMed dataset. **(a)** Distribution of the abstract length in characters. For the distribution of the abstract length over the embedding, see Figure S11. Papers with abstracts shorter than 250 characters were filtered out (see Methods). **(b)** Distribution of the abstract length in words. The smooth peaks visible in panel (a) likely originate from the sharp peaks visible in panel (b), as the journals often specify the maximal allowed abstract length in words (e.g. 150, 200, or 250 words). **(c)** The total number of papers per year, the number of available first/last authors’ first names per year, and the number of inferred first/last author genders per year. The amount of available first names increased dramatically after 2003, when PubMed began incorporating more detailed author information into their database (97.4% of available first names are post-2003).

**Figure S2:**
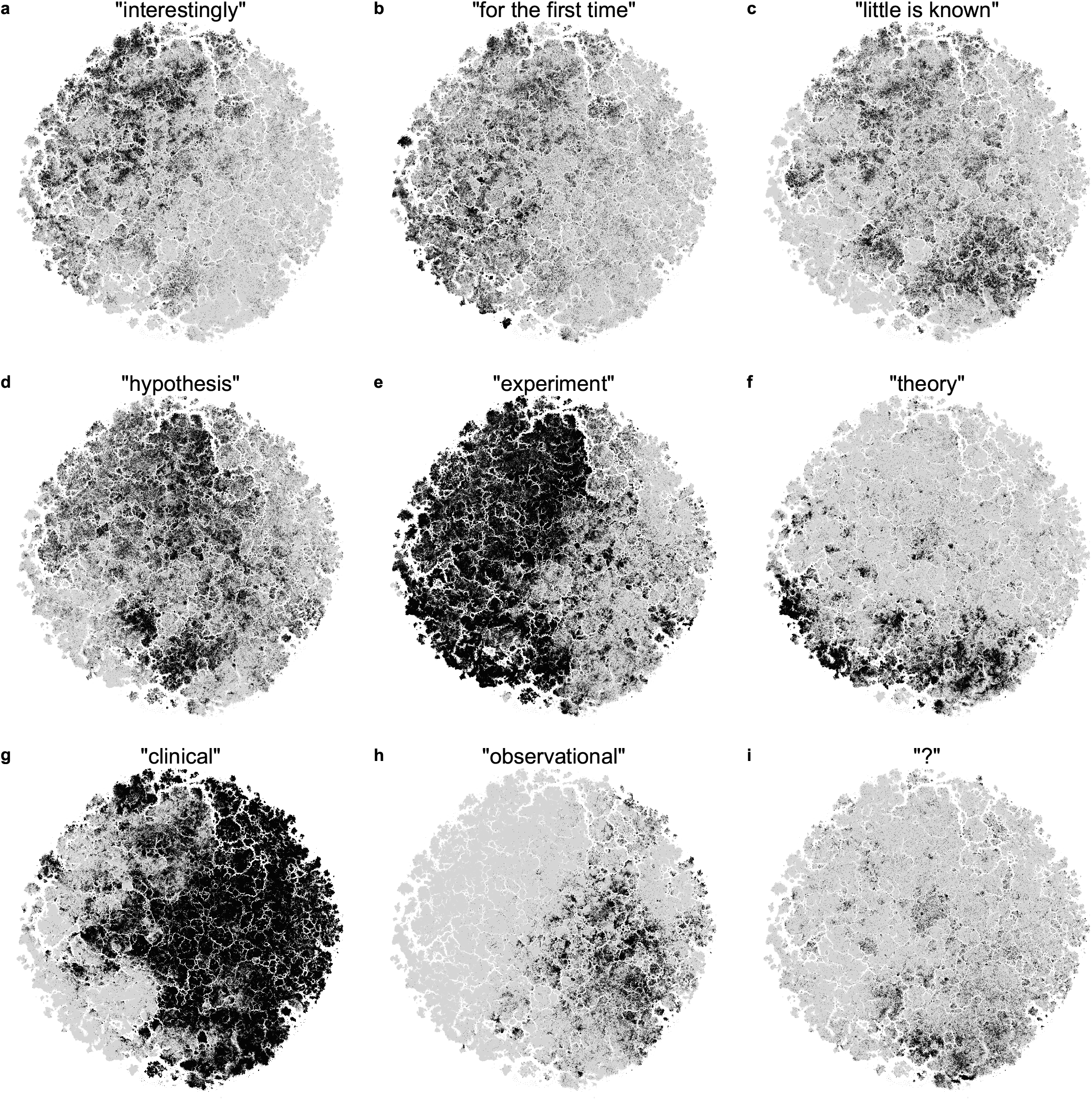
Distribution of some terms and phrases across the biomedical literature. All panels show the embedding based on the PubMedBERT representation, highlighting papers containing particular terms in their abstracts. **(a)** ‘interestingly’, **(b)** ‘for the first time’. Two black islands stand out in the periphery of the embedding: the one in the bottom contains articles reporting new species (‘species nova’) and the one on the left contains articles reporting novel chemical compounds isolated from living organisms. **(c)** ‘little is known’, **(d)** ‘hypothesis’, **(e)** ‘experiment’, **(f)** ‘theory’, **(g)** ‘clinical’, **(h)** ‘observational’, **(i)** ‘?’ (question mark).

**Figure S3:**
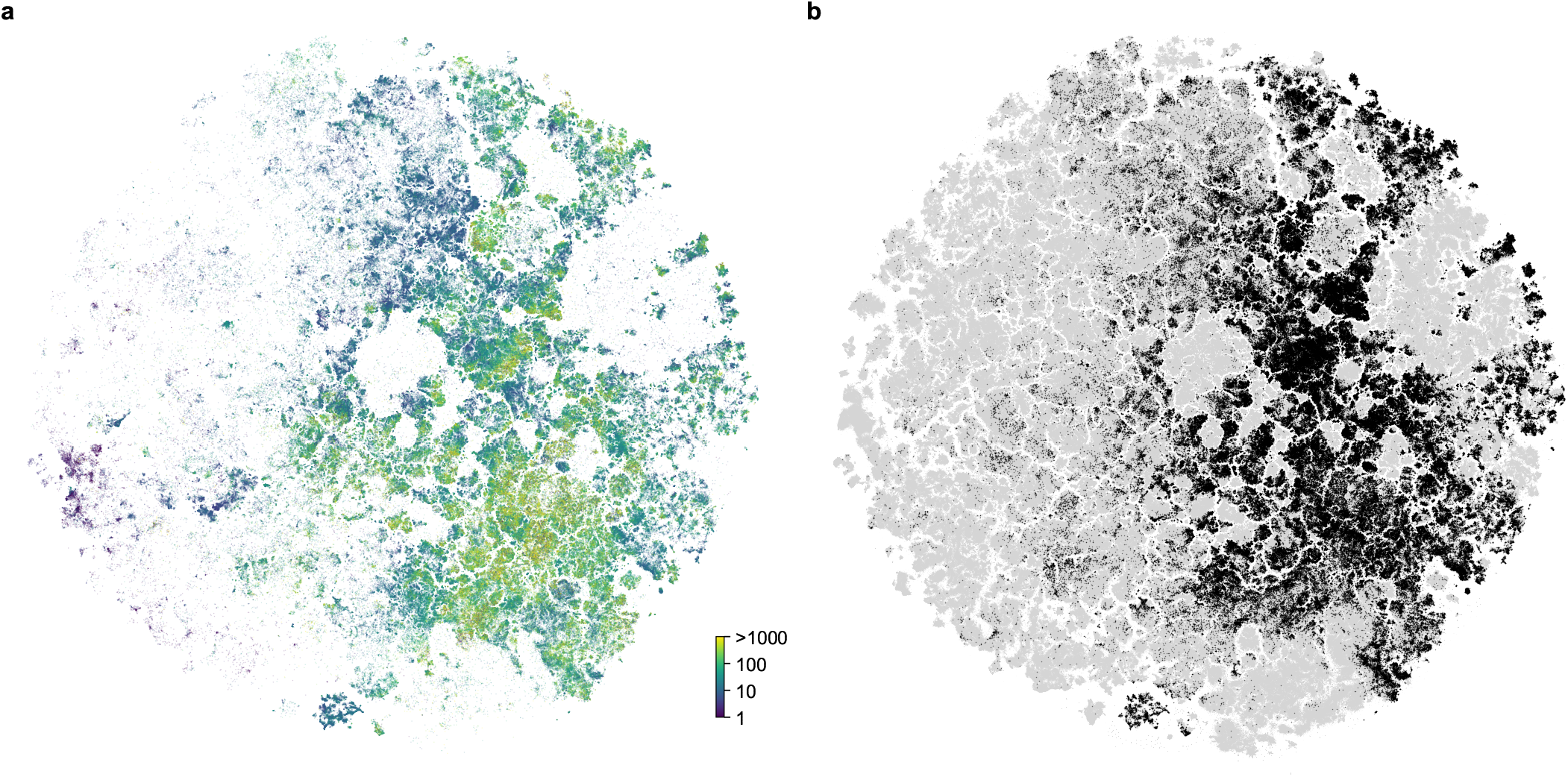
Distribution of reported sample sizes and *p*-values across the biomedical landscape. **(a)** Embedding colored by the sample size reported in the abstract. We used the regular expression `n\s?=\s?(\d+)` to extract the reported sample sizes. If an abstract contained several reported sample sizes, we took the first one. Color scale on the log scale, dark: *n* = 1; light: *n* ≥ 1000. Papers that did not contain this regular expression in their abstract are not displayed. **(b)** Papers reporting *p*-values in their abstracts (containing `‘p=‘` or `‘p<‘` strings, with or without space after ‘p’) are shown in black.

**Figure S4:**
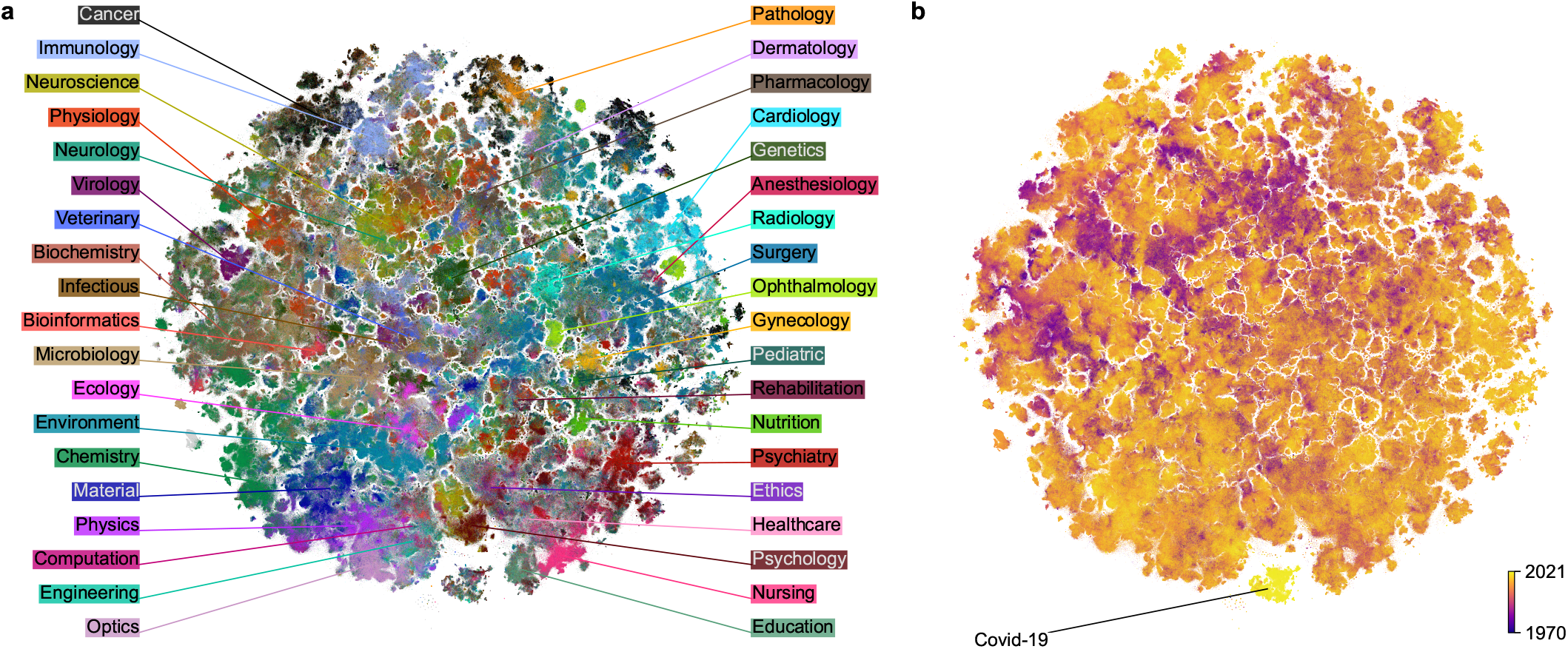
2D embedding based on the TF-IDF representation of the PubMed dataset. **(a)** Colored using labels based on journal titles. Unlabeled papers are shown in gray and are displayed in the background. The TF-IDF-based embedding was flipped to orient it similarly to the BERT-based embedding (Figure 1). **(b)** Colored by publication year (dark: 1970 and earlier; light: 2021).

**Figure S5:**
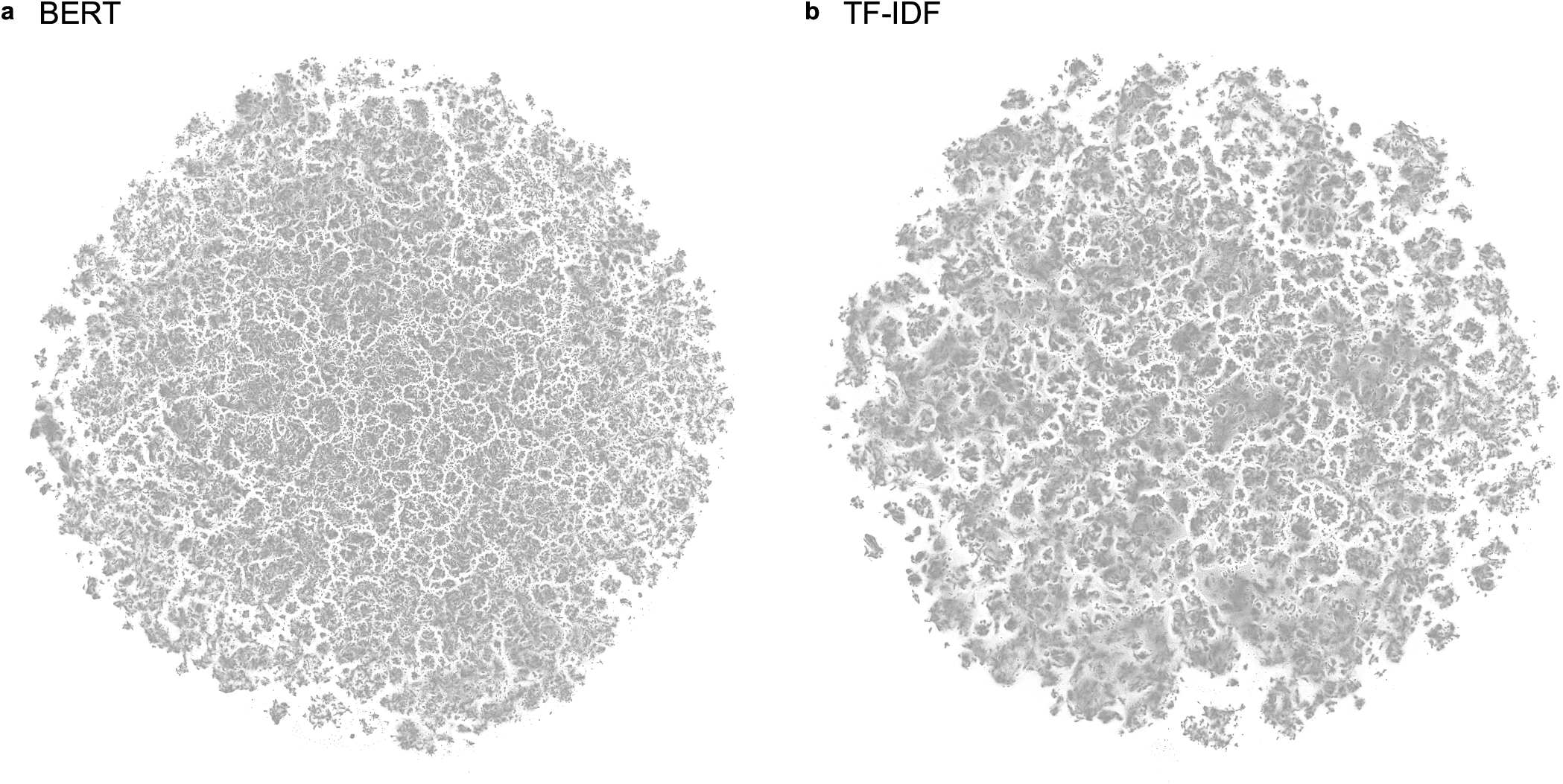
Fine cluster structure in the PubMed embeddings. All points shown in gray to emphasize the cluster structure. **(a)** The embedding based on the PubMedBERT representation. **(b)** The embedding based on the TF-IDF representation.

**Figure S6:**
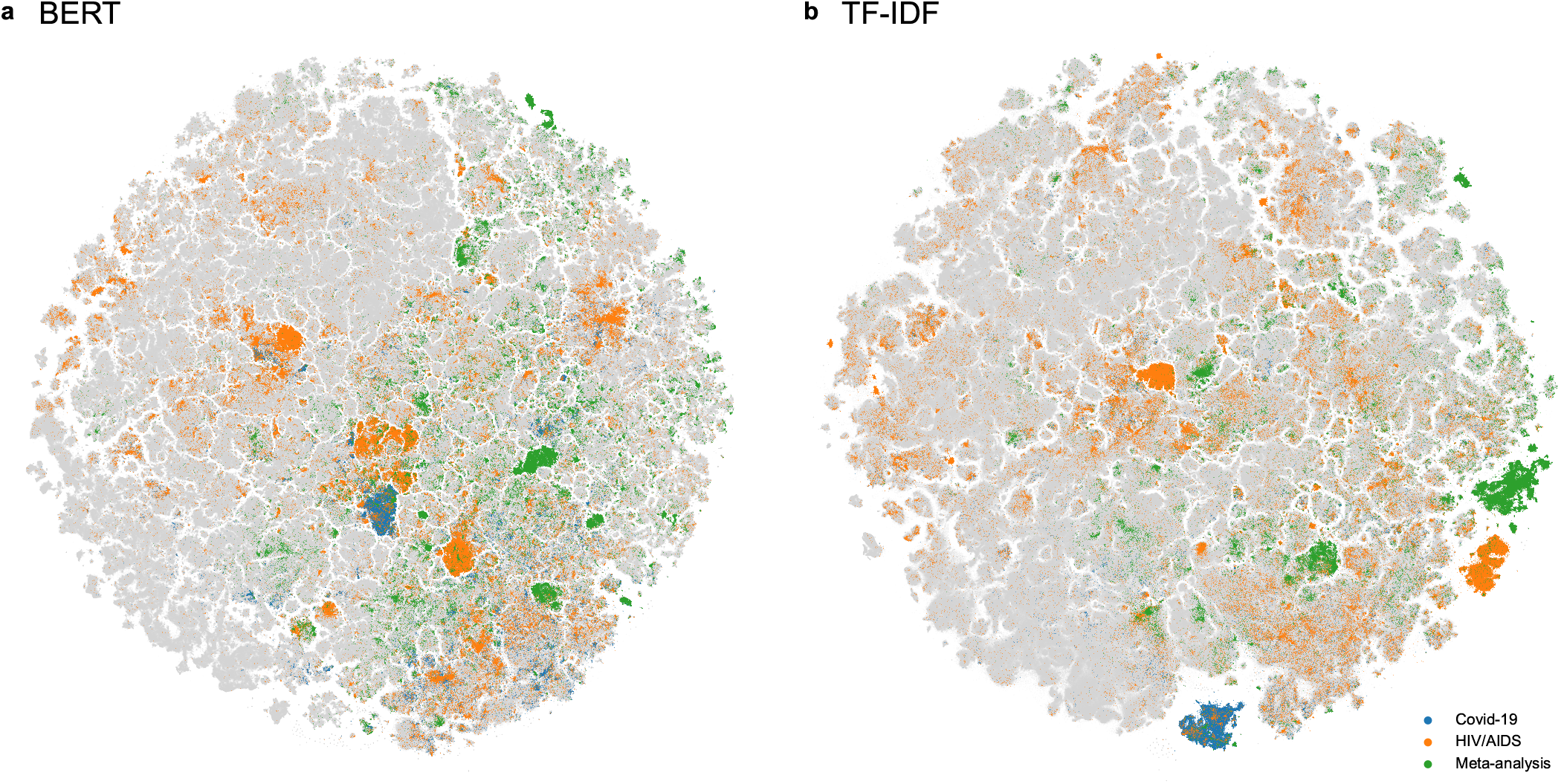
Isolated subcorpora in the PubMed embeddings. Three sets of papers analyzed in Table 2 (Covid-19, HIV/AIDS, meta-analysis) highlighted in both embeddings. **(a)** PubMedBERT-based embedding. **(b)** TF-IDF-based embedding. In the TF-IDF-based embedding, the Covid cluster appeared more separated from the rest of the embedding, and included a larger fraction of Covid papers (86.7%), compared to the BERT-based embedding. Similarly, meta-analysis papers and HIV papers were grouped together and isolated stronger than in the BERT-based embedding. This suggests that TF-IDF representation is more sensitive to the presence of specific keywords than the BERT representation, which is more faithful to semantic similarity between fields (e.g. between Covid papers and the literature on other respiratory diseases).

**Figure S7:**
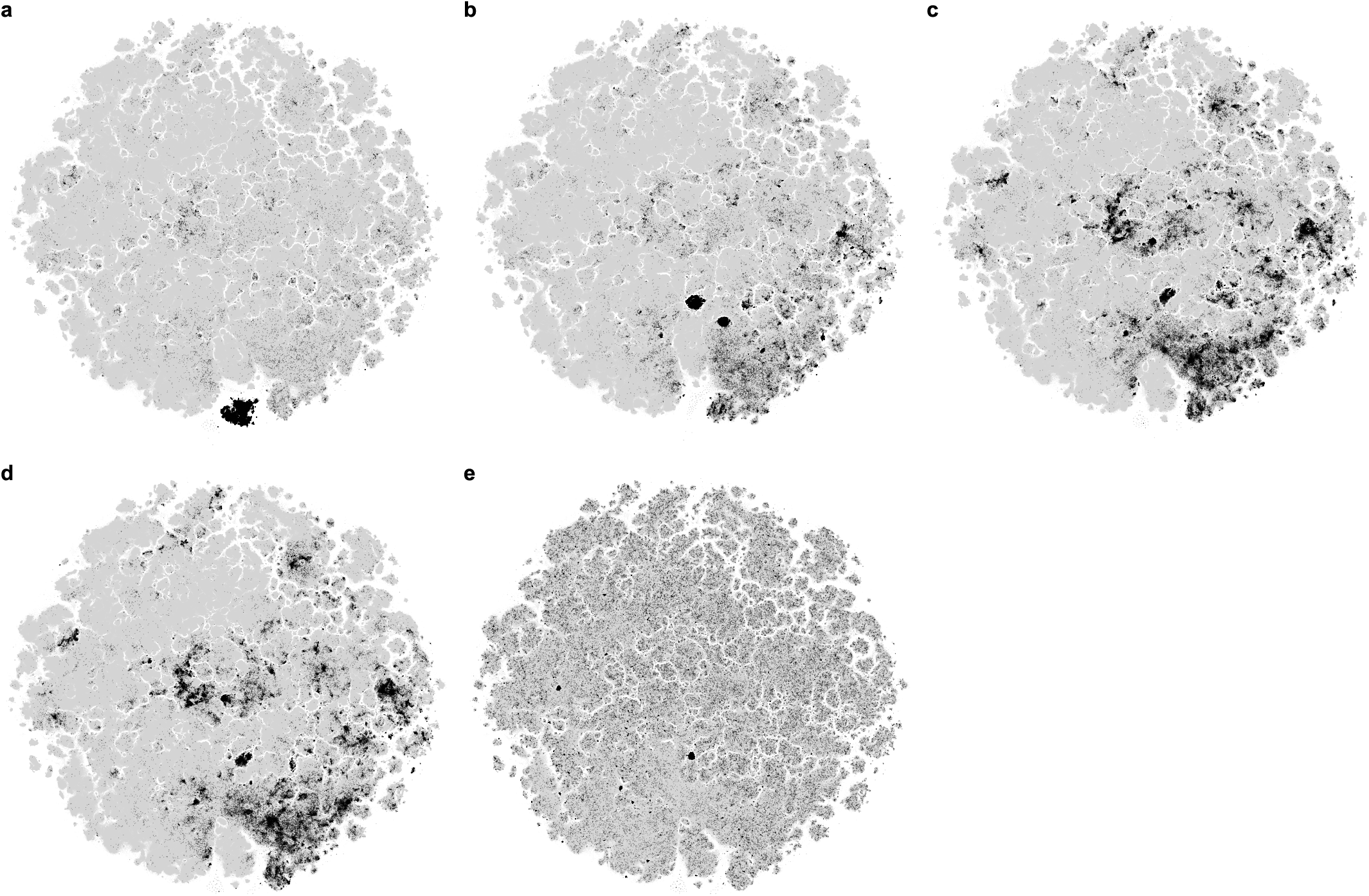
Covid-19 island ablation experiment. The TF-IDF data were altered in several ways and the different versions were used to obtain *t*-SNE embeddings (following SVD and using the same pipeline as described in the Methods). Papers highlighted in black are Covid papers, unless otherwise stated; non-Covid papers are shown in grey in the background. **(a) Unaltered TF-IDF**. The same embedding as in Figure S4. **(b) TF-IDF without the ‘covid’ feature**. We eliminated the feature corresponding to the word ‘covid’ from the TF-IDF data. The Covid island disappeared but the Covid papers were still grouped together in two clear clusters. **(c) TF-IDF without Covid-related features**. We eliminated several Covid-related features (‘covid’, ‘19’, ‘coronavirus’, ‘sars’, ‘cov’, and ‘2019’) from the TF-IDF data. Here the Covid papers were spread out more. **(d–e) TF-IDF with Covid-related features shuffled**. We shuffled the rows of the TF-IDF submatrix with six columns corresponding to the Covid-related features listed above. The other columns were kept intact. This procedure assigns Covid-related features to random papers in the dataset. In the resulting embedding, the Covid papers were spread out similarly to panel (c). In panel (e) we show the same embedding but highlight the random papers that got assigned the Covid-related features after shuffling. These papers are spread out much more uniformly compared to the Covid papers in panel (d), meaning that the actual Covid papers had much more in common (in the TF-IDF space) than only the Covid keywords.

**Figure S8:**
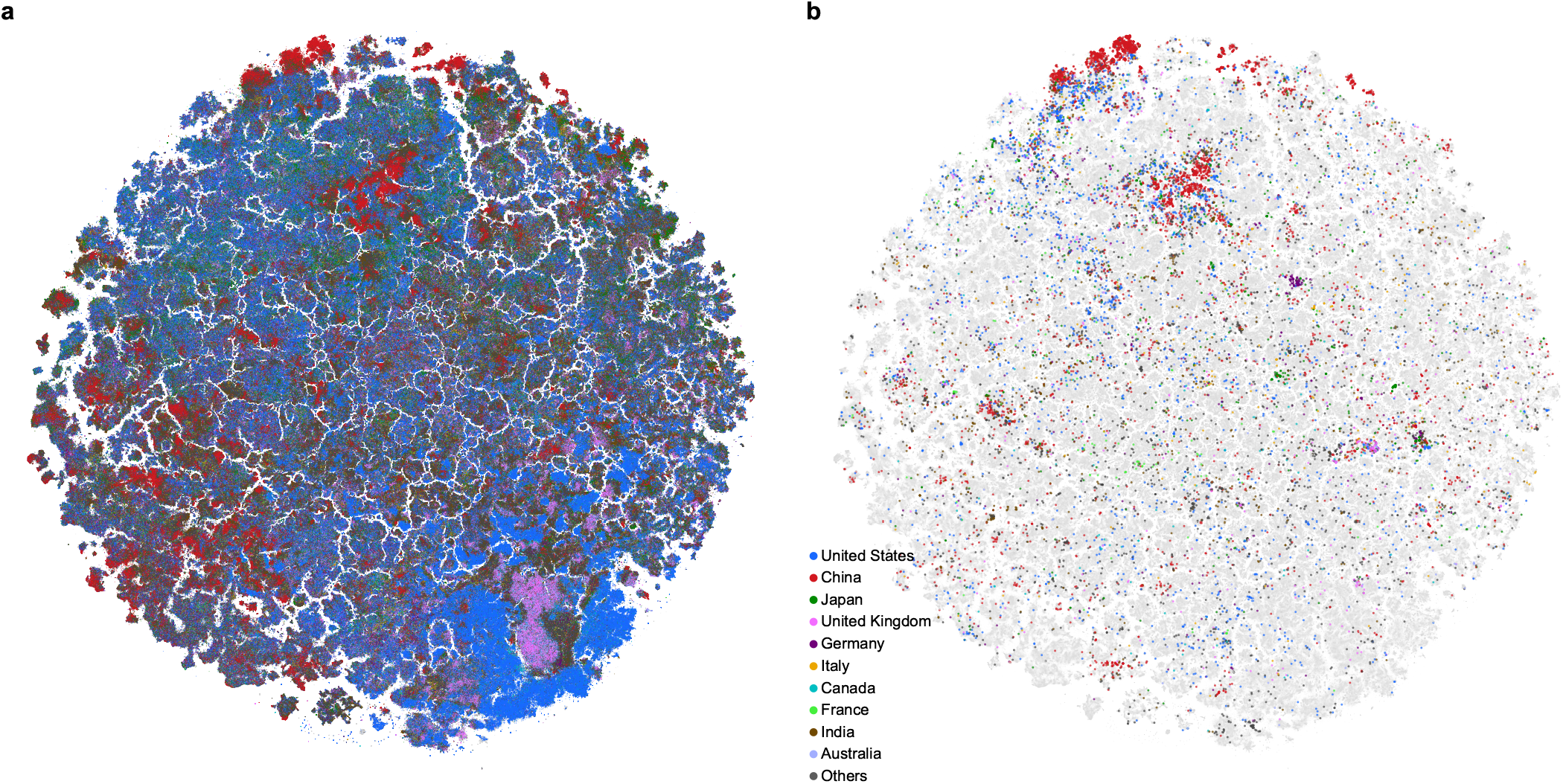
Distribution of affiliation countries across the biomedical landscape. **(a)** Embedding colored by affiliation country of the first author. Top 10 countries by the total number of papers are shown in colors, the rest are shown in dark gray. Papers without country information are shown in light gray and displayed in the background. **(b)** Embedding showing retracted papers (as in Figure 6), colored by their affiliation countries. Several clusters of retracted papers stem mostly from a single country. In some cases, these clusters correspond to a single author involved in a large-scale scientific misconduct (e.g. German cluster: Joachim Boldt; Japanese cluster: Yoshihiro Sato; US cluster: Scott Reuben). In other cases, clusters contain a larger number of papers from many different authors but one single country, which could be an indicator of paper mill activity. Visible separation between the UK and the US papers in social disciplines (lower-right corner) and in some medical islands may be due to differences between the British and the American spelling.

**Figure S9:**
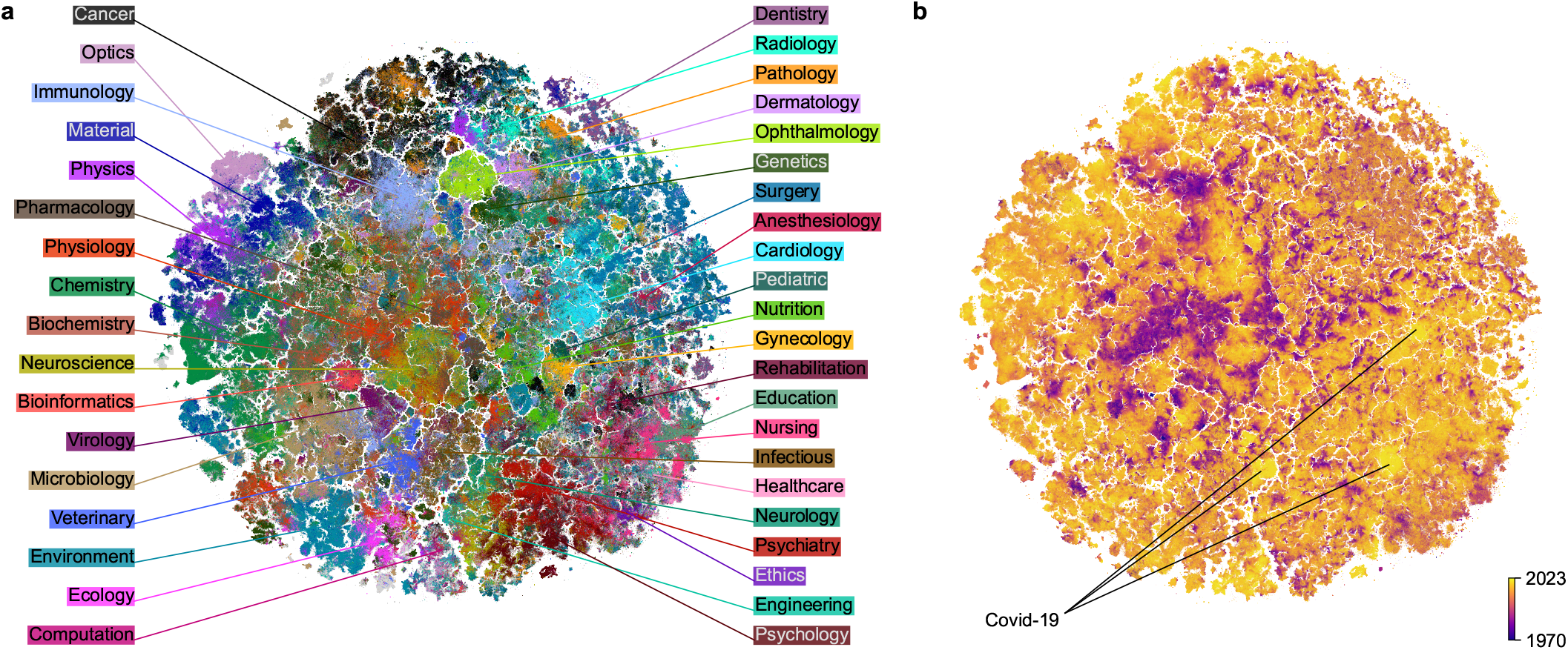
2D embedding of the updated PubMed dataset. Here we used annual PubMed snapshot that included 2022 and 2023 papers, which were not part of the initial dataset shown in Figure 1. Sample size 23.4 M. **(a)** Colored using labels based on journal titles. Unlabeled papers are shown in gray and are displayed in the background. ‘Dentistry’ was added as a new label, not present in Figure 1. **(b)** Colored by publication year (dark: 1970 and earlier; light: 2023). The Covid-19 island present in Figure 1 got now divided into three main clusters: the leftmost island contained articles on epidemiology and vaccinology, the lower right island focused on the societal aspects of the pandemic, and the upper right island focused on clinical and medical aspects of Covid-19.

**Figure S10:**
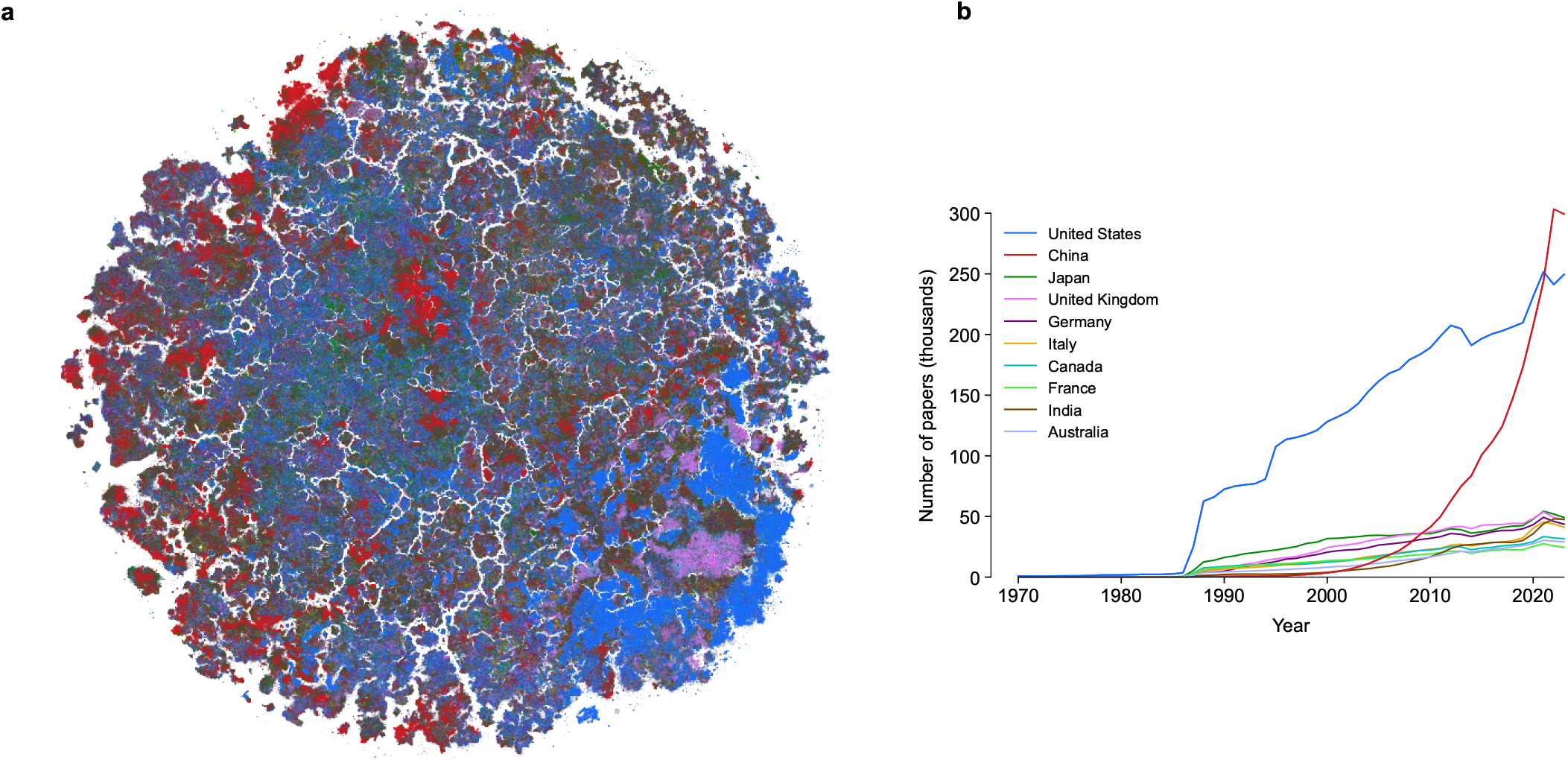
Affiliation countries in the updated PubMed dataset. **(a)** Updated embedding (Figure S9) colored by the affiliation country of the first author. Top 10 countries by the total number of papers are shown in colors, the rest are shown in dark gray. Papers without country information are shown in light gray and displayed in the background. **(b)** The number of papers from each of the top 10 countries over the years. The number of publications from Chinese institutions has increased exponentially over the last years (Else and Van Noorden, 2021), growing from 3.6 K (1.0% of all publications) in 2000 to 303 K (22.2%) in 2022 and surpassing the US in 2021. Note that the 2023 data in this PubMed snapshot were still incomplete.

**Figure S11:**
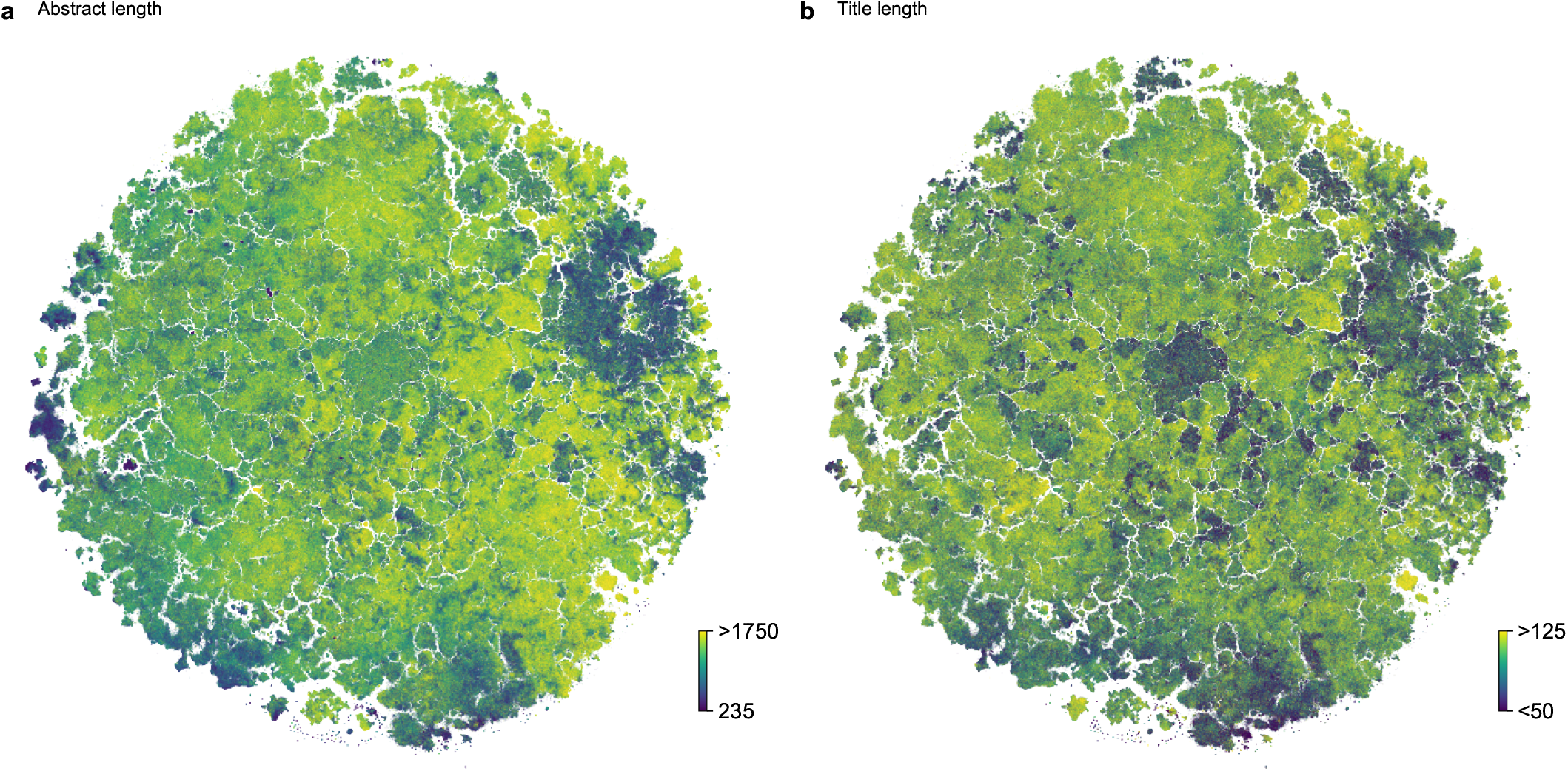
Distribution of abstract and title lengths across the biomedical literature. Both panels show the embedding based on the PubMedBERT representation of the PubMed dataset. **(a)** Colored by the length of the abstract (dark: 235 characters; light: 1750 characters or more). **(b)** Colored by the length of the title (dark: 50 characters or less; light: 125 characters or more).

**Figure S12:**
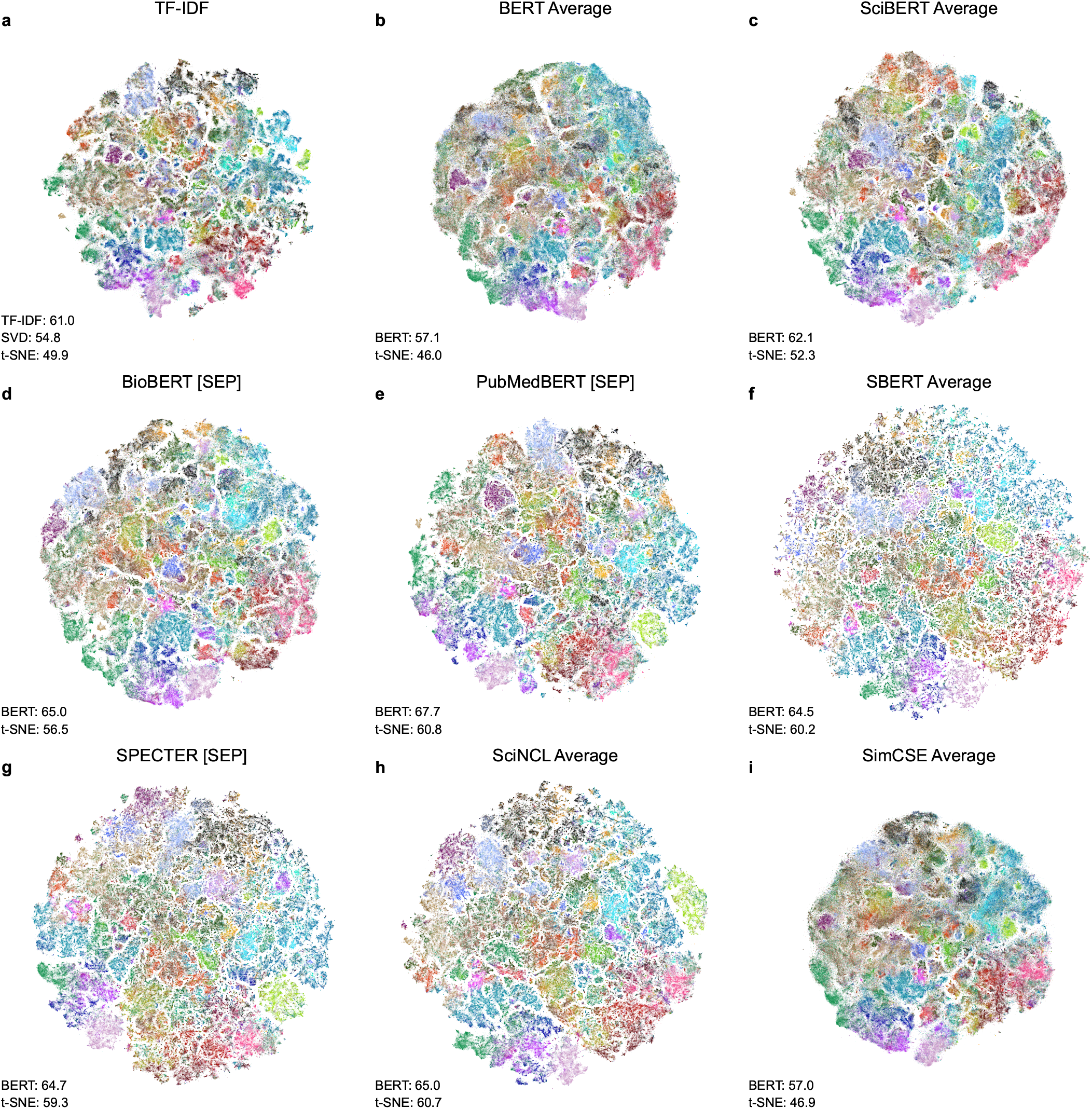
*t*-SNE embeddings of a subset of the PubMed dataset based on different representations. Subset size: 1,000,000 labeled papers. For each BERT-based model, we chose the two-dimensional embedding based on the representation (average, `[CLS]`, or `[SEP]` token) with the highest *k*NN accuracy, see Table 4. The *k*NN accuracies for the high-dimensional and two-dimensional representations are shown in the corner of each panel. The embeddings were flipped to orient them similarly to the embedding of the full dataset (Figure 1). **(a)** TF-IDF (using SVD), **(b)** BERT, **(c)** SciBERT, **(d)** BioBERT, **(e)** PubMedBERT, **(f)** SBERT, **(g)** SPECTER, **(h)** SciNCL, **(i)** SimCSE.

**Figure S13:**
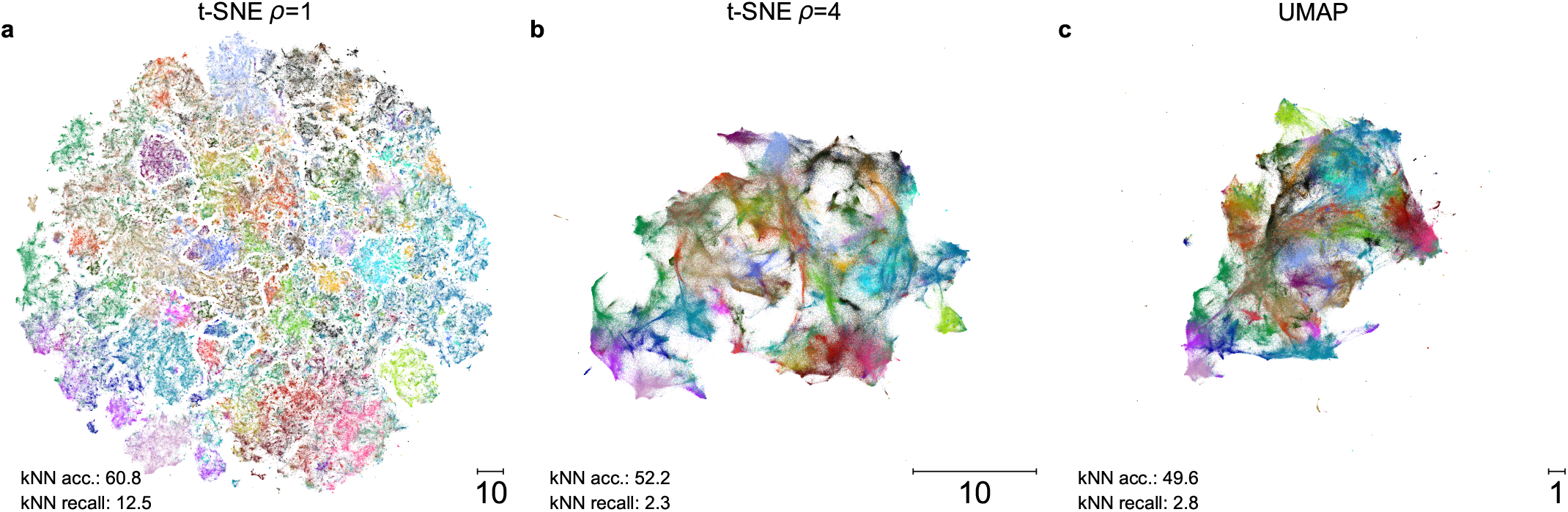
Embeddings of a subset of the PubMed dataset using different neighbor embedding methods. Subset size: 1,000,000 labeled papers. The embeddings were flipped to orient them similarly to the embedding of the full dataset (Figure 1). **(a)** *t*-SNE without exaggeration (*ρ* = 1). **(b)** *t*-SNE with exaggeration *ρ* = 4. **(c)** UMAP.

